# A metabolic atlas of mouse aging

**DOI:** 10.1101/2024.05.04.592445

**Authors:** Steven E Pilley, Dominik Awad, Djakim Latumalea, Edgar Esparza, Li Zhang, Xuanyi Shi, Maximilian Unfried, Shuo Wang, Racheal Mulondo, Sriraksha Bharadwaj Kashyap, Darius Moaddeli, Peter Sajjakulnukit, Damien Sutton, Harrison Wong, Aeowynn J. Coakley, Gilberto Garcia, Ryo Higuchi-Sanabria, Sophia Liu, Bingfei Yu, William B Tu, Brian K. Kennedy, Costas A Lyssiotis, Peter J Mullen

## Abstract

Humans are living longer, but this is accompanied by an increased incidence of age-related chronic diseases. Many of these diseases are influenced by age-associated metabolic dysregulation, but how metabolism changes in multiple organs during aging in males and females is not known. Answering this could reveal new mechanisms of aging and age-targeted therapeutics. In this study, we describe how metabolism changes in 12 organs in male and female mice at 5 different ages. Organs show distinct patterns of metabolic aging that are affected by sex differently. Hydroxyproline shows the most consistent change across the dataset, decreasing with age in 11 out of 12 organs investigated. We also developed a metabolic aging clock that predicts biological age and identified alpha-ketoglutarate, previously shown to extend lifespan in mice, as a key predictor of age. Our results reveal fundamental insights into the aging process and identify new therapeutic targets to maintain organ health.

## Introduction

The average human lifespan worldwide has increased from 66.8 in 2000 to a peak of 73.4 in 2019 (World Health Organization), but this has not been accompanied by a rise in healthspan – how long people live in generally good health. The incidence of age-related chronic diseases, such as cancer, cardiovascular disease and neurodegeneration^1^ is also increasing, fueling a need to uncover drivers of aging and strategies to increase healthy longevity. Recent studies characterizing age-dependent changes at the molecular level^2,3^ have associated aging with a decline of tissue structure and function. These and other studies have identified 12 hallmarks of aging, with several having direct links to metabolic processes, such as epigenetic alterations, loss of proteostasis, deregulated nutrient sensing, and mitochondrial dysfunction^4^. Most strategies that increase lifespan also directly impact metabolism. Examples include: genetic deletion of growth hormone receptor in mice^5^; caloric restriction^1^; dietary methionine restriction^6^; dietary supplementation of alpha-ketoglutarate (AKG)^7^; and mTOR inhibitors such as rapamycin^8^. Conversely, premature aging occurs in some inherited genetic disorders that affect metabolism. Certain types of cutis laxa result from deficiencies in proline metabolism, and progeroid Ehler-Danlos syndrome may be caused by defects in glycosaminoglycan synthesis^9^. The importance of metabolic homeostasis during aging is also highlighted by its association with aging-associated diseases including cancer^10–12^, obesity^13^ and cardiovascular disease^14^. For example, increased levels of circulating methylmalonic acid in adults over 65 years can accelerate cancer development^15^. Dysregulated metabolism is also linked to age-related neurodegeneration, for example induced neurons from Alzheimer’s disease patients have increased expression of PKM2 that causes a reprograming of glycolysis^16^. Combined, these studies highlight the importance of metabolic homeostasis in the aging process, but to date no study has described the metabolic changes that occur throughout life in multiple organs.

Previous studies of age-related changes have been performed in select organs^17–19^, or at other stages of life, such as during fetal development^20^. This contrasts with studies describing other omics of aging across multiple tissues, such as changes in cell composition and gene expression in the Tabula Muris Senis dataset^2^.

Sex-dependent differences in longevity are observed in many species, with females living longer than males in humans^21^. Sex also impacts disease metabolism^22^. In diabetic kidney disease, sex-dependent differences in metabolism enable female kidneys to cope with high glucose concentrations better than males to prevent disease development^23^. Multiple factors including the environment, sex hormones, and sex chromosomes could influence these metabolic differences. Therefore, identifying sex-independent and sex-dependent changes in metabolism during aging is critical to understand why: 1. women live longer than men; and 2. women and men have different disease susceptibility.

Studies also show that lifestyle and disease can uncouple biological age from chronological age, determined primarily using epigenetic aging clocks based on patterns of DNA methylation^24,25^. These clocks also show that biological age could potentially be reversed by interventions such as exercise^26^. However, epigenetic clocks were not developed to identify therapeutic targets or mechanisms of aging. Clocks built on downstream biological processes such as changes in levels of lipids^27^, proteins^28^, RNA^29,30^ or metabolites, could provide these mechanistic and therapeutic insights.

In this study, we described how metabolism changes in 12 mouse organs at 5 ages in both male and female mice. We discovered different patterns of metabolic aging between organs and sexes, identified hydroxyproline as a new marker of aging, and developed a metabolic aging clock based on plasma metabolite levels. This work also provides a resource for other studies investigating age- and sex-specific processes and disease.

## Results

### Overview of the targeted metabolomics data

To profile how organ metabolism changes throughout the murine lifespan, we performed targeted metabolomics on 12 organs from 5 different ages of mice: adolescent (1 month old); young adult (3 months old); and old adult (18 months, 21 months, and 24 months). Each age group contained 7 male and 7 female mice, enabling us to compare age- and sex-dependent changes in metabolism. We matched the mouse strain, ages and diet with the Tabula Muris Senis study^2^, allowing for integration of the two datasets. Within 10 minutes of euthanasia, we harvested the plasma, heart, pancreas, liver, kidney, brain, spleen, quadriceps muscle, lung, thymus, tongue and bladder (Figure 1A). We extracted polar metabolites from the 840 samples and quantified metabolite levels using liquid chromatography with tandem targeted mass spectrometry (LC-MS/MS) using hydrophilic interaction liquid chromatography (HILIC) and dynamic multiple reaction monitoring (dMRM) in negative mode.

**Figure 1:**
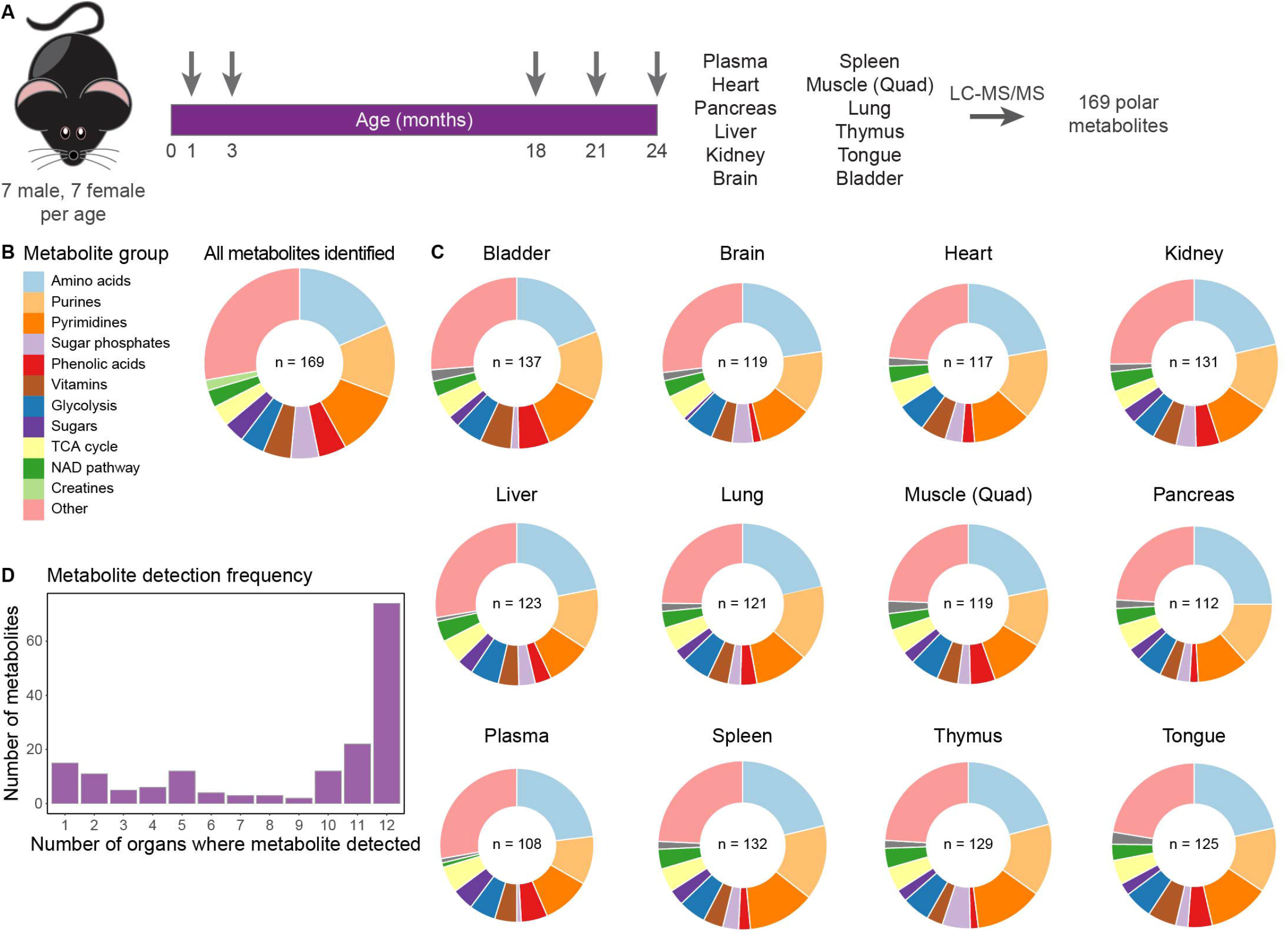
Overview of the Metabolic Atlas of Aging. (A) Schematic of the experimental design. The 5 arrows represent the ages at harvest. (B) Distribution of metabolite classes identified in our targeted analysis. (C) Distribution of metabolite classes identified in each organ. The diameter of each donut is proportional to the number of metabolites (n) identified per organ. Colors indicate the distribution of metabolite groups. (D) The number of organs in which a metabolite is found.

We used Skyline^31^ to analyze the chromatograms and identified metabolites based on overlapping qualified and quantifier peaks and proximity to previously established retention times. Data were normalized to total ion count and log-transformed. Across the dataset, we detected 169 metabolites, covering central carbon metabolism, amino acids, nucleotides and cofactors (Figure 1B). We detected the most metabolites in the bladder (137) and the fewest in the plasma (108) (Figure 1C). Almost half of all the metabolites detected (74) were found in every organ and over 60% were identified in at least 10 organs, whereas 15 metabolites were only found in a single organ (Figure 1D). Similar numbers of metabolites from each metabolite group were identified across all organs, although there were some differences. For example, no sugar phosphates were found in the plasma, and no sugars were found in the heart (Figure S1). We then performed Spearman correlations to determine similarities between samples within and between organs. Samples within individual organs correlated and clustered with each other (Figure 2A), providing confidence in our analysis. Strong correlation within organs across different age groups suggests that metabolic differences between organs are maintained throughout life. Samples from the thymus showed the lowest intra-organ average correlation (0.77), suggesting variability with age, whereas plasma samples showed the highest intra-organ correlation (0.97) (Figure S2). There were also positive correlations between organs, indicating similarities in their metabolic environments. The highest average inter-organ correlation was between the thymus and spleen (0.46), potentially reflecting that these are both lymphoid organs containing cells with similar metabolic requirements. The second highest inter-organ average correlation was between two skeletal muscles, the tongue and the quadriceps muscle (0.34). This may reflect the similarity in their organ function and cellular composition. Of note, the most negative average inter-organ correlation was between the plasma and heart, with the next five most negative all including the plasma, suggesting that the contamination of the organ samples with plasma was minimal, and that the metabolic microenvironment of the organs is distinct from the plasma.

**Figure 2:**
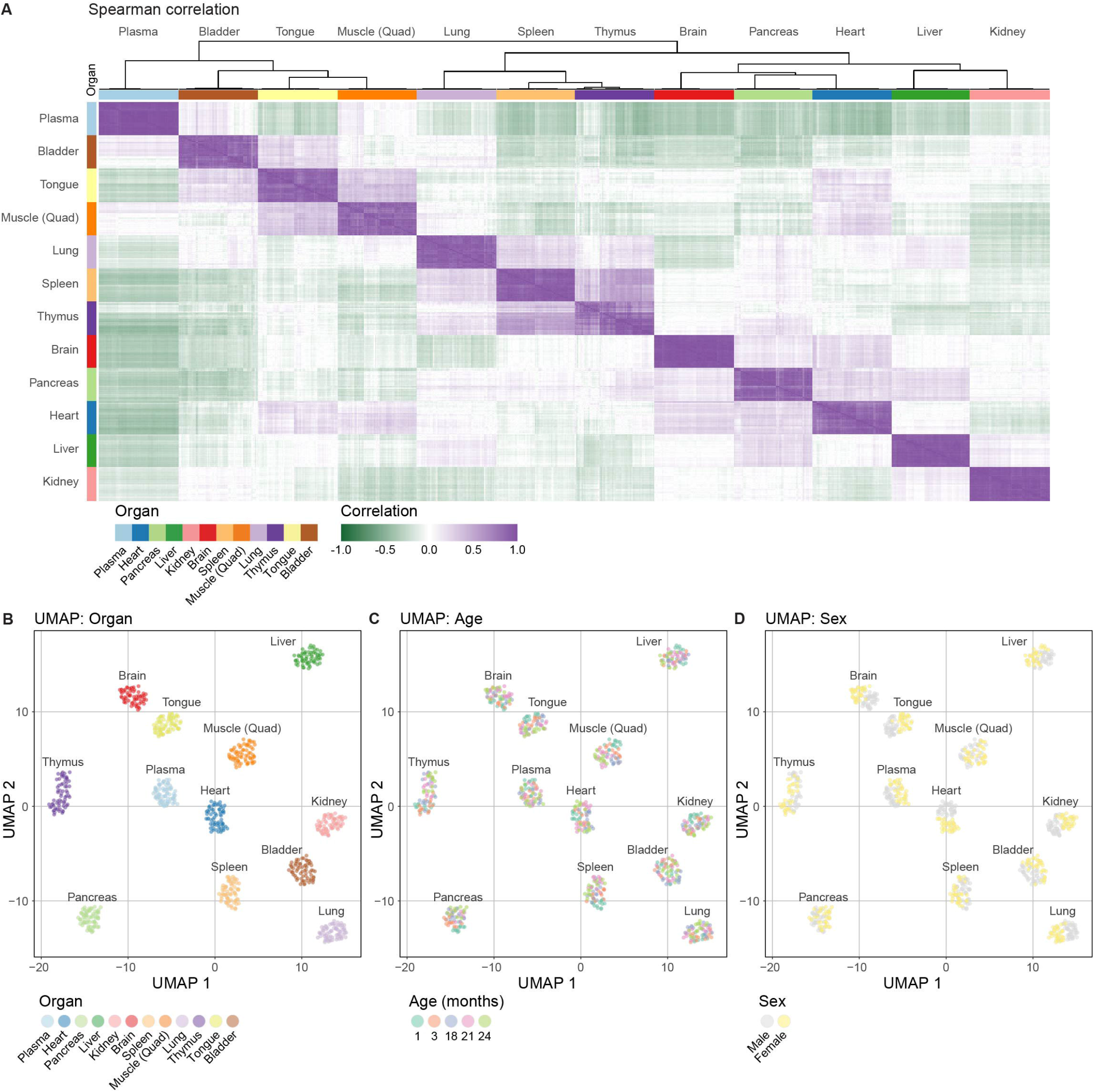
Samples within organs show strong correlation. (A) Unsupervised clustering of all samples based on the Spearman correlation between every sample. (B) UMAP of the data labeled by organ. (C) UMAP of the data labeled by age. (D) UMAP of the data labeled by sex.

Consistent with Spearman correlations, samples within organs also clustered together using Uniform Manifold Approximation and Projection (UMAP) analysis (Figure 2B). We then clustered the samples by age, revealing distinct patterns between organs. The quadriceps muscle, tongue, thymus, spleen and pancreas show clear clustering of samples with age (Figure 2C), whereas other organs – the brain, plasma, heart, kidney and lung – show clearer clustering by sex (Figure 2D).

### Organs show distinct patterns of metabolic aging

Our dataset reveals distinct patterns of metabolic changes during aging in each organ (Figure 3A). Every organ shows age-dependent changes in metabolite levels, although the numbers and trajectories of changes vary between organs (Figures 3A and 3B). For example, 45.7% of detected metabolites show at least one significant fold change during thymus aging, compared to only 7.6% in the brain. A previous study analyzed age-related metabolic changes in 10 different regions of the brain^17^, and found significant metabolic heterogeneity across brain regions. Our brain region agnostic extraction method possibly accounts for fewer significant changes observed.

**Figure 3:**
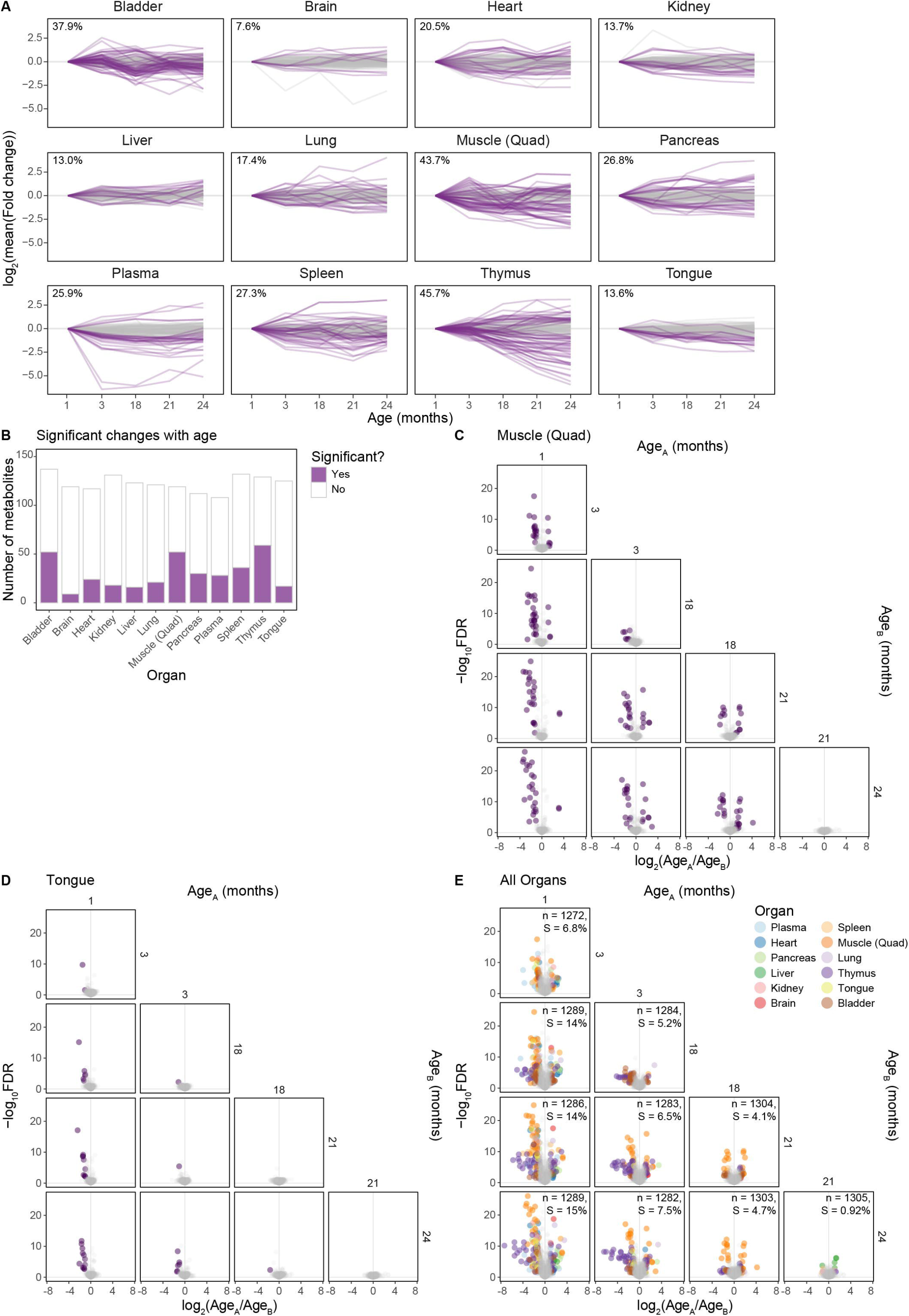
The effect of aging on organ metabolism. (A) Trajectory of metabolite levels in each organ. Each line shows how the levels of an individual metabolite change at each age. All levels are log_2_ mean fold change relative to 1 month. Purple lines represent metabolites that show at least one significant change with age (FDR < 0.05, log_2_ mean fold change < −1 or > 1), gray lines represent metabolites which show no significant changes with age. The percentage of metabolites that show at least one significant change is also shown for each organ. (B) Number of significant metabolites per organ. Bars show the total number of metabolites identified in each organ. Purple indicates the number of metabolites that show at least one significant change with age (FDR < 0.05, log_2_ mean fold change < −1 or > 1). White indicates metabolites that show no significant changes with age. (C) Volcano plot showing the distribution of metabolite changes between ages in the quadriceps muscle. Purple dots represent significant changes (FDR < 0.05, log_2_ mean fold change < −1 or > 1). (D) Volcano plots showing the distribution of metabolite changes between ages in the tongue. Purple dots represent significant changes (FDR < 0.05, log_2_ mean fold change < −1 or > 1). (E) Volcano plots showing the distribution of metabolite changes between ages in all organs. Number of comparisons (n) and percentage of comparisons which are significant (S) shown. Purple dots represent significant changes (FDR < 0.05, log_2_ mean fold change < −1 or > 1).

Although there was a positive correlation between samples in the quadriceps muscle and the tongue (Figure S2), these two organs showed different metabolic changes during aging (Figure 3A). Almost 44% of quadriceps muscle metabolites showed at least one significant fold change in levels between ages, whereas only 14% of metabolites were significantly altered in the tongue (Figures 3C and 3D). This difference may reflect the age-related decline in muscle mass and function observed in the quadriceps but not in the tongue^32^.

Reduced metabolic function is associated with aging^33^, and our dataset supports this by revealing that more metabolites decrease with age than increase. For example, 56% of 5,136 metabolites are reduced in older mice compared to 1 month old mice (Figure S3A). 646 of those comparisons are significant (FDR < 0.05, log2 fold change > |1|), and 72% of those are a decrease from 1-month-old mice (Figure S3B). Interestingly, only the liver shows a general increase in the levels of metabolites with age (Figure S3C).

The most metabolically distinct age group is the adolescent 1 month old mice, which shows the most significant differences when compared to all other groups (Figure 3E). Since 1-month-old mice have not reached adulthood, metabolic differences associated with hormone levels and growth are expected. Despite this, there are more significant changes between 3-month-old mice and 24-month-old mice (96 significant changes out of 1282 comparisons) than between 1 month old mice and 3-month-old mice (87 significant changes out of 1272 comparisons), indicating that the metabolic changes imposed by old age are as drastic as those experienced between adolescence and adulthood (Figure 3E). Metabolism continues to change into old age: there are almost as many significant differences between mice aged 18 months and 24 months (61) as between mice aged 3 to 18 months (64).

### Metabolic aging is different between the thymus and spleen

The thymus and the spleen are lymphoid organs and show the highest inter-organ average correlation (Figure S2). However, they are known to be affected differently by age: the thymus undergoes thymic involution, reduced mass and decreased cellularity beginning in early adulthood, whereas the spleen maintains mass in mice^34^ and only slightly decreases in size in humans^35^. We therefore asked whether there were age-related differences in metabolism related to these differences. Consistent with the large changes in its structure and function, the thymus showed the greatest change in metabolite levels with age of all organs we profiled, with almost 50% of metabolites showing age-dependent significant changes (Figure 3A). PCA showed that PC1 accounts for 45% of the variation in the thymus, indicating that age is the predominant contributor to the variation between samples (Figure 4A). 1- and 3-month-old thymus samples overlapped and clustered separately to 18-, 21- and 24-month-old thymuses, indicating that most age-related changes in the thymus occur in adulthood between 3 and 18 months. In contrast, PCA of spleens showed 1-month-old mice clustering separately from the other ages, indicating that the greatest age-related change in spleens occurs between 1 and 3 months (Figure 4B). This suggests that the transition between adolescence and adulthood in the spleen is associated with more metabolic changes than the transition into old age.

**Figure 4:**
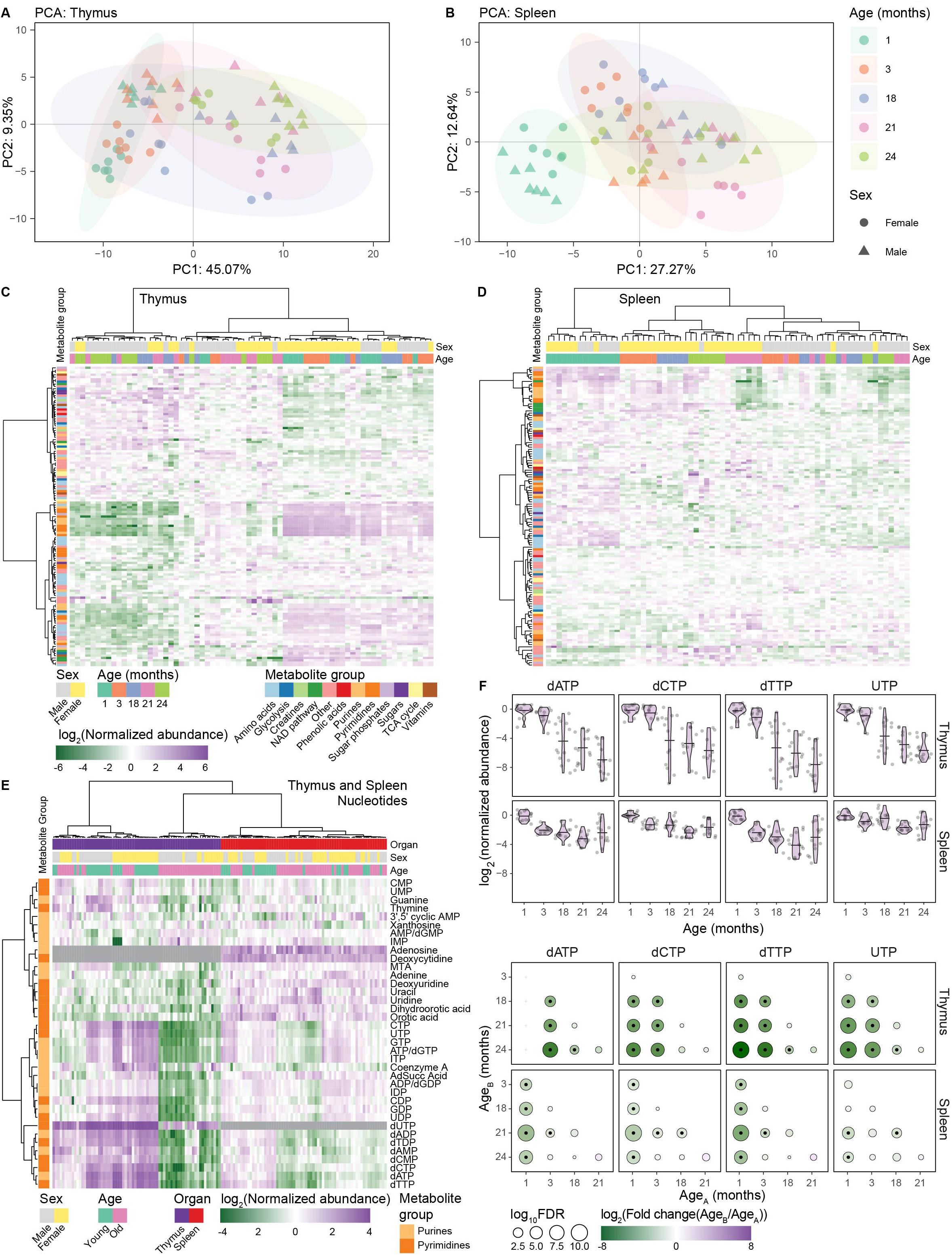
Metabolic aging is different between the thymus and spleen. (A) Principal component analysis of thymus samples. Shape distinguishes sex and color distinguishes age. (B) Principal component analysis of spleen samples. Shape distinguishes sex and color distinguishes age. (C) Heatmaps showing the pareto scaled log_2_ transformed abundance of thymus metabolites and samples sorted by unsupervised clustering. Samples are sorted along the x axis with their age and sex indicated and metabolites are sorted in the y axis with their group indicated. (D) Heatmaps showing the pareto scaled log_2_ transformed abundance of spleen metabolites and samples sorted by unsupervised clustering. Samples are sorted along the x axis with their age and sex indicated and metabolites are sorted in the y axis with their group indicated. Legend as in (C). (E) Heatmap of purine and pyrimidine nucleotide levels in the thymus and spleen. ‘Young’ represents 1- and 3-month-old mice, and ‘old’ represents 18-, 21- and 24-month-old mice. Samples are sorted along the x axis with their age and sex indicated and metabolites are sorted in the y axis with their group indicated. Dark gray squares on heatmap indicate metabolite not detected. 5’-methylthioadenosine (MTA), adenylosuccinic acid (AdSucc acid). (F) Top. Violin plots showing the log_2_ transformed normalized (to 1-month old) abundance of selected nucleotides in the thymus and spleen. Horizontal lines indicate the mean log_2_ transformed normalized abundances at each age. Bottom. Significance plots of nucleotide changes. Circle diameter shows the log_10_ FDR for the indicated metabolites for each comparison of age B relative to age A. Circle color indicates the size of the log_2_ transformed fold change. A black dot in the center of the circle indicates the comparison has an FDR < 0.05 and a log_2_ mean fold change of < −1 or > 1. A gray cross indicates the comparison was excluded as the data was found to not be normally distributed.

We then visualized all metabolic changes in thymuses (Figure 4C) and spleens (Figure 4D) and observed that there are more and larger changes in metabolite levels in the thymus compared to the spleen. Spleens from all the 1-month-old mice clustered together, whereas the other ages are clustered first by sex and then by age, suggesting male and female spleens undergo different metabolic changes after adolescence (Figure 4D). As shown by the PCA, thymuses from the younger mice (1- and 3-month-old) clustered separately from the older 21- and 24-month mice (Figure 4C). However, thymuses from the 18-old-month mice showed sex specific differences in metabolic aging. The majority of female 18-month-old mouse thymuses clustered with the younger mice, whereas most male 18-month-old mouse thymuses clustered with the older mice, suggesting that there is earlier thymic metabolic aging in male mice than female mice. Previous studies show that thymic involution occurs earlier in males than in females, and our metabolic phenotype could be reflective of this difference in the rate of thymic involution^36,37^. To investigate metabolic links to thymic involution, we next identified metabolites that show age-related changes during aging. One cluster of 15 metabolites showed a large age-dependent decrease, and 13 of those metabolites are nucleotides (Figure 4C). We then clustered the thymuses and spleens based only on nucleotide levels (Figure 4E) and observed that nucleotide levels decrease with age more in the thymus than in the spleen. Representative nucleotides highlight this difference (Figure 4F). This age-associated decrease in nucleotide levels in the thymus correlates with age-associated thymic involution, which is characterized by a reduction in size, loss of thymic epithelial cells and reduced production of naïve T-cells^38^. Female mice maintain thymus nucleotide levels at 18 months more than male mice (Figure S4A), matching with previous data showing thymic involution happens faster in males^36^. In humans, thymic involution begins early in childhood and continues through adulthood^39^, but this transition may occur more gradually in mice^38^. This suggests that the age-associated drop in nucleotides is reflecting reduced proliferation, which is associated with aging in thymuses^40–42^. To test this, we analyzed data from the Tabula Muris Senis, a previously published study of RNA expression in single cells during aging^2^ and identified that the proportion of rearranged double positive thymocytes dropped with age in the thymus, including a large drop from 21- to 24-month-old mice^2^ (Figures S4B, S4C, and S4D). This indicates that with age, thymuses produce a less diverse repertoire of T-cell receptors, which would reduce their ability to target non-self antigens. Furthermore, we analyzed spatial RNA-sequencing data^43^ and found that a proliferation gene expression signature drops with age in multiple immune compartments (Figure S4E). Whilst there are peaks in the proliferation signature before 1-month-old (∼30 days), the decreases carry on beyond 3 months (∼100 days) into old age. This correlates with the drop in thymus nucleotide levels observed at 18 months and older. Taken together, the metabolic and RNA expression data showed that major biological changes occur with age in the thymuses of mice, which reflects the dynamics of thymic involution.

### Sex impacts organ metabolism

Our study can also identify age-independent sex differences in metabolism, as it included organs from 7 male and 7 female mice at each age (Figure 1A). To give an overview of sex-dependent differences in metabolism at each age, we ranked the ratios of metabolite abundances between males and females at each age in each organ (Figure 5A). This analysis revealed different patterns of sex-specific changes in metabolism between organs. The heart, thymus, quadriceps muscle and bladder showed many significant sex differences at all ages whereas the pancreas showed sex differences at specific ages, with more metabolites affected by sex in 1-month-old mice than at other ages (Figure S5). Other organs showed metabolic divergence between the sexes during aging, as evidenced by 1-month-old mice clustering together in the spleen (Figure 4B) and kidney (Figure S6A) and then separating at 3-months-old and older.

**Figure 5:**
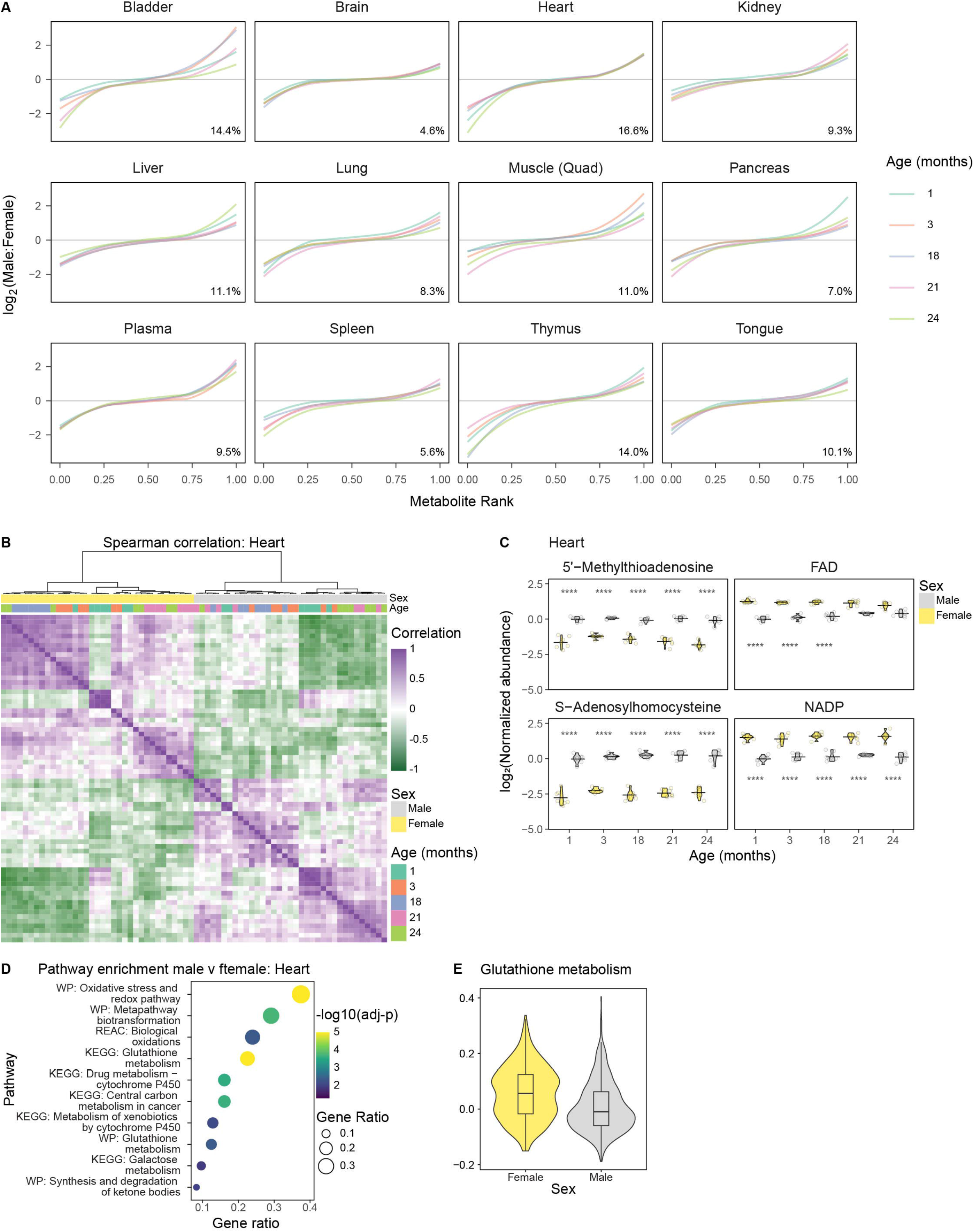
The effect of sex on organ metabolism. (A) Sex-specific differences in heart metabolites at each age. Each line shows the log_2_ ratio of metabolite levels in males to females at a different age. Metabolites are ordered along the x-axis by the size of the ratio at that age in that organ. The percentage in the bottom right corner indicates the percentage of comparisons in that organ which are significant (FDR < 0.05, log_2_(male:female) < −1 or > 1). (B) Unsupervised clustering of heart samples by Spearman correlation between each sample. (C) Selected metabolic differences between male and female hearts. Violin plots showing the log_2_ transformed normalized (to 1-month old) abundance of the indicated metabolites. Horizontal lines indicate the mean log_2_ transformed normalized abundances at each age. **** indicates FDR < 0.0001 for male to female comparisons. (D) Pathway enrichment of gene expression differences between 18-month-old male and female hearts. Data analyzed from the Tabula Muris Senis^2^. (E) Pathway enrichment of the significant differential glutathione metabolism genes between male and female hearts. Data analyzed from the Tabula Muris Senis^2^.

The heart showed the highest percentage of significant differences between males and females of the same age, and these sex differences were maintained throughout life. This is reflected in the Spearman correlation of the heart samples, which cluster into separate male and female groups (Figure 5B). We also observed similar clustering by sex in PCA (Figure S6B). These sex dependent differences in the heart correlated with sex-dependent differences in heart tissue cell composition, with female mice having more fibroblasts and fewer endothelial cells than male mice (Figure S6C). Within the male and female clusters, there is weak clustering by age, and this may reflect that the levels of several metabolites are consistent through life but different in males and females. To answer this, we identified metabolites that are consistently different between males and females at all ages. Interestingly, 5’-methylthioadenosine (MTA) and S-adenosylhomocysteine (SAH) – which are both involved in methionine metabolism – are higher in males than females throughout life (Figure 5C). In the cell, methionine is converted into the methyl donor S-adenosylmethionine (SAM) using ATP. Upon donation of its methyl group, SAM is converted into SAH. SAM is also required for polyamine synthesis. MTA is also produced by these reactions and is converted back into methionine by the methionine salvage pathway^44^ (Figure S6D). Methionine restriction affects male and female energy metabolism differently, as in young males it causes fat loss and retention of lean muscle, but young females preserve fat and lose lean muscle^45^. Our results suggest that there are differences in methionine metabolism in male and female hearts throughout life. Despite the differences in MTA and SAH levels, methionine levels are similar in males and females (Figure S6E). There are also significant differences in the levels of MTA and SAM between males and females in the bladder, lung, thymus and tongue (Figure S6F). Three out of four of these organs have higher levels of MTA and SAM in males.

Two other heart metabolites that showed consistent sex differences across age were FAD and NADP^+^ (Figure 5C). To determine whether differences in FAD and NADP^+^ levels were reflective of wider differences in oxidation state in male and female hearts, we looked at the levels of other redox associated metabolites. NADPH is significantly higher in younger female hearts (Figure S6G) and the overall NADP^+^:NADPH ratio is higher in older females than in older males (Figure S6H). However, there are no significant differences between the NAD^+^:NADH ratio in male and female hearts at any age (Figure S6I). Interestingly, female hearts have a higher ratio of reduced to oxidized glutathione (GSH:GSSG) compared to male hearts (Figure S6J). This correlates with pathway analysis on mouse heart RNA expression levels, which shows that oxidation and glutathione pathways are enriched in female hearts^2^ (Figures 5D and 5E). These metabolic changes correlate with previously identified sex differences in redox regulation in the heart and may contribute to increased levels of cardiovascular disease in men^46^.

### Trans-4-Hydroxyproline depletion is a marker of organ aging

We next used our dataset to identify metabolites that show the largest changes during aging. Using a regression analysis, we calculated the rate of change in metabolite levels with age (Figure 6A). The top 15 steepest decreases with age were all found in the thymus and 13 of these were nucleotides, confirming our observation in Figure 4E. The metabolite that showed the steepest increase with age is itaconic acid in the lung. The spleen also showed a similar increase in Itaconic acid. Macrophages produce itaconic acid in response to infection to reduce inflammation^47^, and increases in levels with age may therefore reflect age-associated changes in immune system populations and activity.

**Figure 6:**
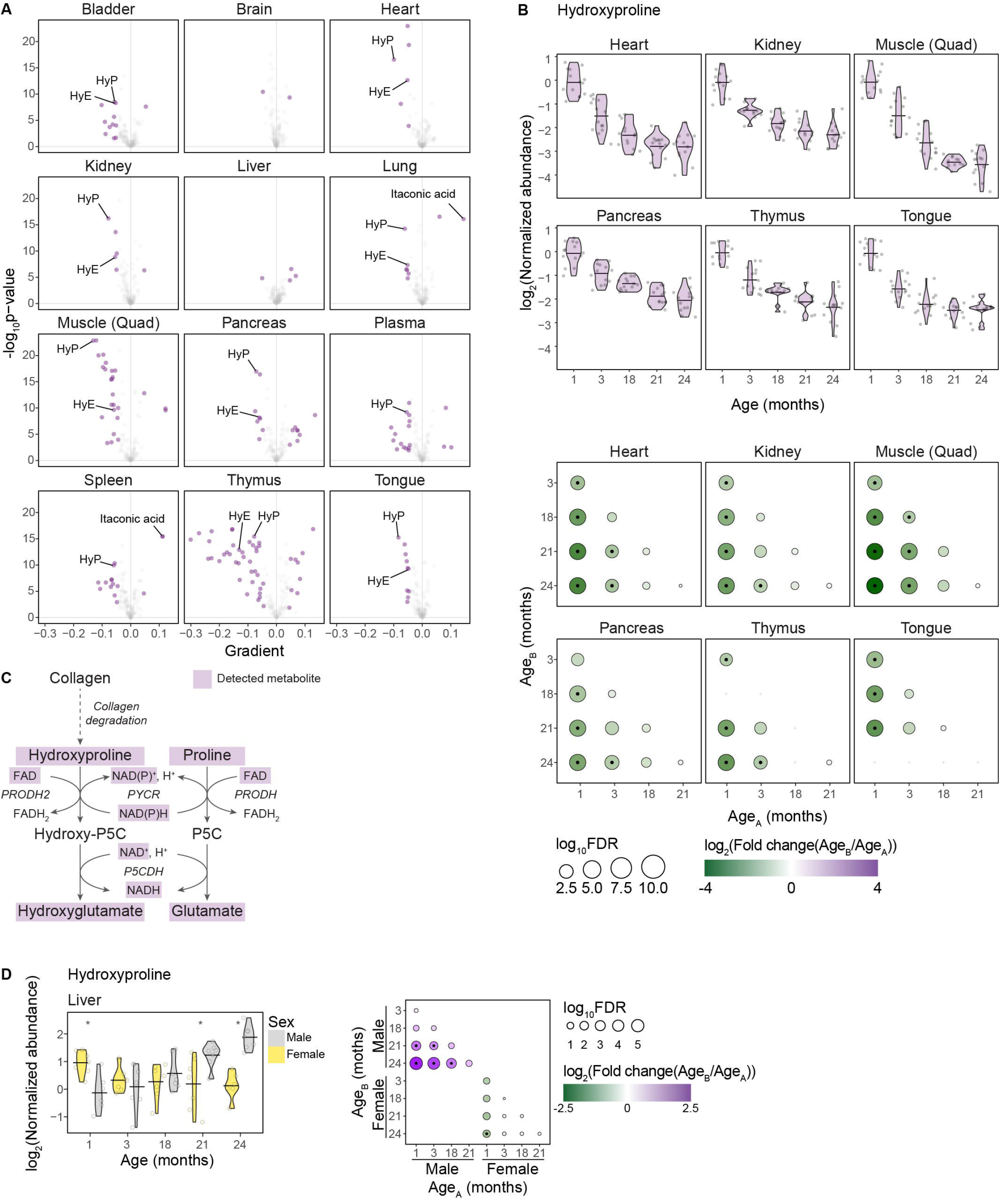
Hydroxyproline shows the most consistent age-dependent change across the data. (A) Volcano plots showing the gradients from regressions of the log_2_ normalized (to 1-month old) abundance over age, against the -log_10_(p-value) of that regression, for each organ. Points in purple represent significant regressions (p-value < 0.05, gradient < −0.05 or > 0.05). Hydroxyproline (HyP), hydroxyglutamic acid (HyE) and itaconic acid are labelled. (B) Top. Violin plots showing the log_2_ transformed normalized (to 1-month old) abundance of hydroxyproline in the heart, kidney, quadriceps muscle, pancreas, thymus and tongue. Horizontal lines indicate the mean log_2_ transformed normalized abundances at each age. Bottom. Significance plots of age-related changes in hydroxyproline levels. Circle diameter shows the log_10_ FDR for each comparison of age B relative to age A. Circle color indicates the size of the log_2_ transformed fold change. A black dot in the center of the circle indicates the comparison has an FDR < 0.05 and a log_2_ mean fold change of < −1 or > 1. A gray cross indicates the comparison was excluded as the data was found to not be normally distributed. (C) Schematic of hydroxyproline and proline catabolism. Detected metabolites are in purple boxes. (D) Hydroxyproline levels increase in male livers during aging. Left. Violin plots showing the log_2_ transformed normalized (to 1-month old) abundance of hydroxyproline in the liver separated by sex. Horizontal lines indicate the mean log_2_ transformed normalized abundances at each age for males and females. Right. Significance plots of age-related changes in hydroxyproline levels. Circle diameter shows the log_10_ FDR for each comparison of age B relative to age A. Circle color indicates the size of the log_2_ transformed fold change. A black dot in the center of the circle indicates the comparison has an FDR < 0.05 and a log_2_ mean fold change of < −1 or > 1. A gray cross indicates the comparison was excluded as the data was found to not be normally distributed.

Trans-4-hydroxyproline (hydroxyproline) showed the most consistent change across lifespan. Hydroxyproline levels were reduced in 11 out of 12 organs with age, and 10 out of 12 showed significant negative regressions. Although the regression was not significant in the brain, there was a significant decrease in the level of hydroxyproline between 1- and 24-month-old mice Figure S7A). The organs that showed the steepest decrease in hydroxyproline levels were the quadriceps muscle, tongue and heart (Figure 6B). Hydroxyproline is the hydroxylated form of proline and is only produced by post-translational hydroxylation of proline in proteins, meaning that free hydroxyproline is only produced by protein degradation. The biggest source of free hydroxyproline is collagen^48,49^, where it is essential for collagen stability^50^ (Figure 6C).

Therefore, changes in free hydroxyproline levels may reflect changes in collagen dynamics. Collagen is a key component of the extracellular matrix (ECM), and ECM dysregulation is associated with aging^4,51^. Age-related fibrosis is characterized by the accumulation of ECM components and stiffening of connective tissue^52^. Several hallmarks of aging contribute to dysregulated ECM dynamics, and ECM composition can influence longevity^53^. The reduced levels of free hydroxyproline that we observe may therefore be caused by fibrosis-related reductions in the rate of collagen degradation across organs. We also observed reduced levels of 4-hydroxy-L-glutamic acid (hydroxyglutamic acid) (Figure 6A & S7B), an intermediate in the hydroxyproline degradation pathway^50^ (Figure 6C). Hydroxyglutamic acid is reduced during aging in the same six organs that show the steepest decreases in hydroxyproline (Figure 6A & 6C). The liver was unique in its pattern of age-related changes in hydroxyproline levels: while female livers showed a decrease in hydroxyproline levels consistent with other organs, male livers had increased hydroxyproline levels during aging (Figure 6D). Liver hydroxyglutamic acid levels showed similar trends, with an age-dependent increase in males (Figure S7C). These results indicate that age-dependent changes in collagen dynamics may be different in male livers compared to other organs. Other organs do not show as drastic sex-dependent differences in hydroxyproline or hydroxyglutamic acid levels (Figure S7D & E). The quadriceps muscle showed the strongest reduction in hydroxyproline levels with age. To investigate further, we analyzed the expression of ECM genes in quadriceps muscle in the Tabula Muris Senis database^54^. Matching our metabolite data, we found reduced expression of multiple collagen genes, including *Col1a2*, *Col1a1*, *Col3a1* and *Col6a1* (Figure S8A), matching results from a previous study^55^. We also saw reduced gene signatures of ECM pathways (Figure S8B). Taken together, reduced skeletal muscle collagen expression may therefore account for reduced hydroxyproline levels.

### Creating a metabolic aging clock to predict biological age

Epigenetic clocks use patterns of DNA methylation to predict biological age and age-related pathologies^25^. However, these clocks were not developed to determine mechanisms of aging or identify age-related therapeutic interventions. We therefore used our data to develop a metabolic aging clock that can predict age and identify key metabolites related to aging. To generate the metabolic aging clock, we used plasma metabolite data as plasma is the most accessible material for other studies to use and will therefore be the easiest to translate into humans. We randomly split the data into a training and a testing set and built several models from the data (Figure 7A). We then trained the best performing model on the principal components and the raw data (Figure S9A). We next identified the most important features (metabolites) that enhance the ability of the model to predict age (Figure S9A). In total, 56 mice made up the training set used to build a model (Figure 7B) from 8 different metabolites, each with different coefficients describing their weighting and importance to the model (Figure 7C).

**Figure 7:**
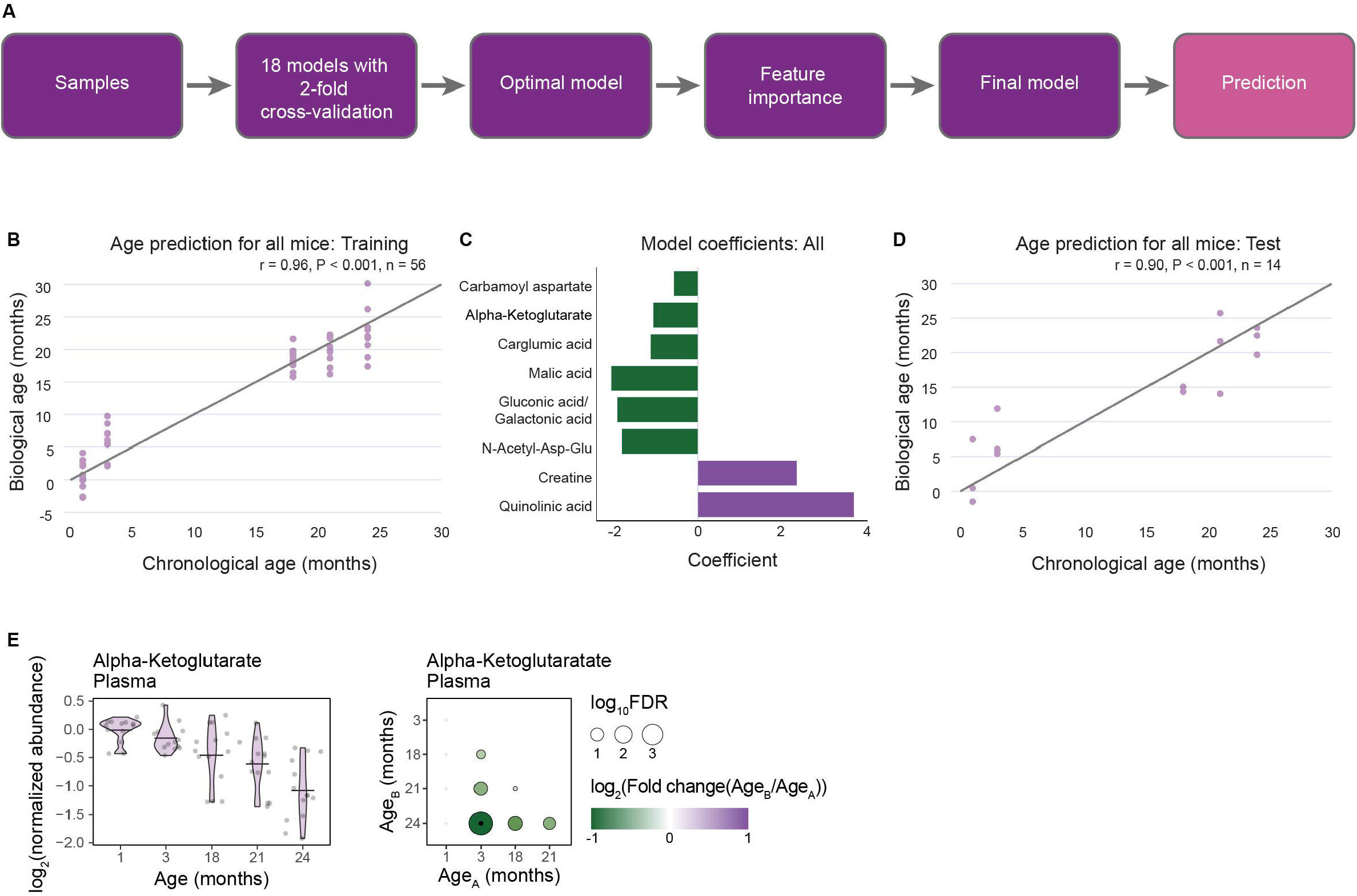
A metabolic aging clock can predict biological age. (A) An overview of the pipeline used to construct the metabolic aging clock. (B) Training set data. The chronological age of samples plotted against the predicted biological age of samples in the training set. ‘r’ indicates Pearson correlation coefficient. (C) Coefficients of the 8 metabolites used by the model to predict age. (D) Test set data. The chronological age of samples plotted against the predicted biological age of samples in the test set. ‘r’ indicates Pearson correlation coefficient. (E) Left. Violin plots showing the log_2_ transformed normalized (to 1-month old) abundance of alpha-ketoglutarate in the plasma. Horizontal lines indicate the mean log_2_ transformed normalized abundances at each age. Right. Significance plots of age-related changes in alpha-ketoglutarate levels. Circle diameter shows the log_10_ FDR for each comparison of age B relative to age A. Circle color indicates the size of the log_2_ transformed fold change. A black dot in the center of the circle indicates the comparison has an FDR < 0.05 and a log_2_ mean fold change of < −1 or > 1. A gray cross indicates the comparison was excluded as the data was found to not be normally distributed.

When tested on the 14 test samples, this model showed a strong ability to predict age, reaching a correlation of 0.90 (Figure 7D). From the 8 metabolites that make up the clock we identified a potential mechanistic driver of aging: alpha-ketoglutarate (AKG) (Figure 7C). We found that serum AKG levels drop with age and are significantly lower in old mice compared to young mice (Figure 7E). Previous studies show that AKG supplementation extends lifespan in several models^7,56^, but no studies have investigated whether AKG levels drop during aging. Our model has therefore identified a metabolite that is reduced during aging, contributes to predicting age and can increase lifespan. Because we saw sex differences in metabolic aging, we also built predictive models of age for males and females individually (Figure S9B, S9C, S9D, and S9E). The male-specific clock used 12 metabolites (Figure S9D), 5 of which were used in the sex-independent model, and predicted age with a strong correlation of 0.93 with the test data (Figure S9F). The female clock was built from only two metabolites (Figure S9E), one of which – quinolinic acid, a product of tryptophan metabolism – was used in both the male model and sex-independent model. The female clock also predicted age with a strong correlation of 0.86 (Figure S9G). These models show that we can predict age from the levels of a select group of plasma metabolites in both a sex-independent and sex-dependent manner.

## Discussion

Here we present an overview of metabolic changes during aging in 12 organs from both male and female mice. Previous studies have focused on single organs^17^, but this atlas represents the first characterization of the metabolism in a broad range of organs and ages in both sexes.

### Thymus metabolism is strongly affected by age

We show that different organs have distinct metabolic phenotypes, although there are also similarities between organs. Interestingly, the plasma shows the lowest degree of similarity with other organs based on Spearman correlation. This indicates that contamination of organ samples with plasma was not significant.

Our data suggests that some organs are better at maintaining metabolic homeostasis during aging than others, although the reasons may vary between organs. For example, the thymus shows the most age-dependent changes of all organs, and this might be related to a decline in organ function with age. Thymic involution occurs gradually throughout life in mice, leading to the loss of T cell development and a reduction in thymus mass. These large changes in organ biology and function would be expected to be accompanied by metabolic changes. Almost 50% of thymus metabolites changed significantly with age, indicating major age-dependent metabolic rewiring. The decrease in nucleotide levels, along with gene expression data suggest the metabolic changes relate to reduced proliferation and thymic involution in older mice.

### Quadriceps muscle metabolism is more affected by aging than the tongue

Our dataset contains two skeletal muscles, the quadriceps and the tongue. Despite the positive correlation between metabolites in the quadriceps and the tongue, there are differences during aging. The tongue shows fewer age-dependent changes in metabolites than the quadriceps.

This is expected since muscle loss during aging is seen in the quadriceps, but to a lesser extent in the tongue^57,58^. It is not known why the tongue is protected from muscle loss, but there is evidence to suggest that tongue strength does not decline during healthy aging^59^. Whether this is connected to the ability of the tongue to maintain metabolic homeostasis is not known, but identifying how this occurs could identify new approaches to reduce age-associated sarcopenia in other muscles.

### Male and female hearts are metabolically distinct at all ages

Our study also uncovers metabolic differences between male and female mice. Some differences are age-dependent, whereas others are age-independent. The heart shows large age-independent sex differences in metabolism, particularly in redox biology. This may contribute to the observed sex differences in age-associated heart dysfunction in humans, and will need to be confirmed in human tissue. Sex differences in the liver and kidney are more age-dependent, only occurring in groups older than 1 month. This matches a previous study showing that there are many sex-dependent differences in kidney gene expression, but these differences only become apparent after 1-2 months of age^60^. A recent study of diabetic kidney disease linked its accelerated progression in males to different metabolic responses to increased circulating levels of glucose in males and females, and that this is caused by increased levels of dihydrotestosterone in males^23^. Why some organs show more differences with age and others show more differences with sex is unclear. It is important to determine which sex differences are due to hormonal differences or genetic differences. For example, males and females have hormone dependent differences in one-carbon metabolism in the liver and responses to methionine restriction^45,61^. Uncovering these relationships between sex-dependent metabolic differences and pathological differences could be used to direct sex-specific therapies.

### Hydroxyproline levels decline during aging

Identifying metabolites that change across multiple organs could uncover new targets to increase the healthspan of those organs. We identified that hydroxyproline levels are reduced in 11 out of 12 organs during aging. Hydroxyproline is a key component of collagen fibers in the ECM, accounting for approximately 13% of their weight^62^. Other components of the ECM have been linked to aging, for example, increasing hyaluronan synthesis can improve healthspan of mice^53^. Altered collagen synthesis, breakdown and structure influence the development of fibrosis, which can affect multiple tissues and has previously been linked with aging^51^. Since collagen is the largest source of free hydroxyproline, it is likely that the observed age-associated decrease in hydroxyproline levels is linked to changes in collagen dynamics: reduced levels of free hydroxyproline may indicate fibrosis. It is also possible that reduced hydroxyproline levels could alter other metabolic pathways during aging. The role of hydroxyproline in normal cellular metabolism is not well characterized, but it is possible that altered levels of hydroxyproline could impact proline metabolism. Hydroxyproline is oxidized by PRODH2, which is analogous to PRODH, the first enzyme in proline catabolism. PRODH activity produces reactive oxygen species^63^ and can act as a tumor suppressor^64^ and contribute to cell survival^65–67^, and PRODH2 may have a similar role^68^. Both hydroxyproline and proline metabolism also use P5CDH and PYCR, suggesting that altered hydroxyproline metabolism could affect proline metabolism^50^. We propose that the age-associated depletion in free hydroxyproline levels can impact the aging process and age-related diseases, although the extent by which this occurs is not known.

### A metabolic aging clock can identify potential drivers of aging

DNA methylation clocks have been used to quantify differences between chronological and biological age. Some epigenetic clocks have also been designed to identify disease and the effects of therapeutics. However, it is not known whether changes in DNA methylation are drivers of aging^4^, or if the methylation status can be used to mechanistically characterize aging. Because metabolites are the downstream readout of cellular processes, an aging clock built on metabolite levels could be used to predict age and offer mechanistic insight. Our metabolic aging clock predicts age from metabolite levels in the plasma with high accuracy, in both sex-independent and sex-dependent formats. Our model selected 8 metabolites for age prediction. One of these, AKG, extended lifespan and healthspan when supplemented in the diet of mice. However, age-dependent changes in AKG levels have not previously been identified. Our model has therefore identified a metabolite that is reduced during aging, contributes to predicting age and can increase lifespan.

In summary, our study uncovers age- and sex-dependent changes in metabolism in 12 organs. These changes reveal fundamental insights into the aging process and identify new therapeutic targets to maintain organ health.

### Study limitations

Our study uses LC-MS/MS based targeted metabolomics to uncover how metabolite pools change during aging in 12 organs in male and female C57BL/6 mice, but it does not address how metabolic flux is altered. Our data suggests that flux studies will uncover further changes in metabolism during aging. Profiling changes in immune cell metabolism during aging could also expand upon data and offer further insight into the differences between thymus and spleen aging.

We focused our study on one mouse strain, C57BL/6NCrl, as it is a commonly used strain, which makes our data relevant to many fields of study. The strain also matches that used in the Tabula Muris Senis, allowing us to incorporate gene expression data into our study. Although we expect many of our observations to be extendable to multiple species, it will still be necessary to characterize how metabolism changes during aging in other systems such as non-human primates and humans.

## Methods

### Animals

We purchased male and virgin female C57BL/6NCrl mice from Charles River (RRID:IMSR_CRL:027) and housed them at USC. Mice were housed under identical conditions at both Charles River and USC (12 hr/12 hr light dark cycle) and fed the same diet ad libitum (NIH-31 diet). The 1-month-old, 3-month-old, and 18-month-old mice were kept at USC for one week before harvesting. The 21-month-old and 24-month-old mice arrived aged 18 months and were aged at USC for the remaining 3 months and 6 months respectively.

### Organ harvest

All tissues were collected beginning at 3 pm on each day of harvest. We used 4% isoflurane to anaesthetize the mice. We then weighed the mice and collected blood via cardiac puncture into EDTA collection tubes and placed them on ice. We then collected organs in the following order: heart, thymus, lungs, liver, pancreas, spleen, kidney, bladder, quadriceps muscle, tongue, and brain. The organs were immediately snap frozen in liquid nitrogen and stored at −80C until metabolite extraction. Organ collection finished by 5 pm. All animal care and procedures were approved by the USC Department of Animal Research. To obtain plasma, we centrifuged EDTA-tubes containing blood at 4°C at 2,000 *g* for 15 minutes. We collected supernatants and added 20 μL to 80 μL 100% LC-grade methanol. For tissue homogenization, we put tissue samples in 10 μL ice-cold 4:1 LC-grade Methanol:Ultrapure-water per mg tissue and homogenized them using a bead homogenizer (Bead Mill 4, Fisherbrand). We centrifuged plasma supernatants and tissue homogenates at 4°C at 21,300 *g* for 10 minutes and collected the metabolite-containing supernatants. We dried down 100 μL of supernatant from each sample (equivalent to 10 mg of tissue) using a vacuum centrifuge and stored them at −80°C until further processing.

### LC-MS/MS

Samples were analyzed as previously described^69^. Dried supernatants reconstituted in 50ul of LC-grade Methanol: Ultrapure-water mix (1:1) and then analyzed using the Agilent 1290 Infinity II LC –6470 Triple Quadrupole (QqQ) tandem mass spectrometer (MS/MS). Compound optimization, calibration and data acquisition was performed using the Agilent MassHunter Workstation Software LC/MS Data Acquisition for 6400 Series Triple Quadrupole MS (software version 10.1) The targeted metabolomics measured 242 metabolites from a reference library in negative mode.

### LC-MS/MS Data processing

We loaded chromatograms into Skyline^31^ (Skyline Daily 23), integrated all peaks and manually curated them for their similarity to the retention time and overlap between qualifier and quantifier peaks. Random batches were analyzed in parallel with Agilent MassHunter Workstation Software LC-MS Data Acquisition for 6400 Series Triple Quadrupole MS with Version 10.1. Curated metabolite abundances were exported for further processing in Microsoft Excel to identify isomers. In R v4.3.1, we normalized peak abundances by the total ion count in each sample and log_2_ transformed them. One sample was removed because of a clerical error. For principal component analysis, we pareto-scaled abundances, and identified and removed two outlier samples (D2, G8) from the data. We normalized abundances to the level at 1-month old and then assessed the normality of the relative abundances of each metabolite in each group using the Shapiro-Wilks test and excluded values with a z-score higher than 2 or lower than −2. We calculated fold-changes and p-values between age (both sex-independent and sex-separate) and sex using the limma package. P-values were adjusted using the false discovery method. Ggplot2 was used for graph construction.

### Thymus Single Cell RNAseq Analysis

The Tabula Muris Senis dataset^2^ was utilized to uncover change in thymus cellular composition during aging. Processed Seurat V5 dataset of all cells (356,213 cells from 23 tissues) was downloaded from https://cellxgene.cziscience.com/collections/0b9d8a04-bb9d-44da-aa27-705bb65b54eb. Cells from thymus tissue were subsetted according to the tissue type embedded in metadata. Cells from donor 24-M-60 were excluded because this sample was dominated by antigen-presenting cells which contained very few thymocytes.

#### Preprocess

After standard Seurat preprocess, canonical correlation analysis (CCA) integration^70^ was performed to eliminate the batch effect between cells from 10X 3’ v2 and Smart-seq2 assays. Ensembl IDs were converted into gene symbols by BiomaRt^71^ R package.

#### Clustering and annotation

After the first-round clustering at a resolution of 0.8, thymocyte clusters were identified according to high expression of Cd3e, Tcf7 and Lck. Thymocytes were subsetted, and the second-round clustering at a resolution of 2 was performed to annotate thymocyte subtypes. These subtypes were identified according to marker gene expression: 1) double negative cells (Cd4-, Cd8a-, Il2ra+); 2) proliferating double positive (DP) cells (Cd4+, Cd8a+, Mki67+); 3) DP cells undergoing rearrangement (Cd4+, Cd8a+, Rag1+); 4) DP cells between rearrangement and selection (Cd4+, Cd8a+, Rag1-, Itm2a-); 5) DP cells undergoing selection (Cd4+, Cd8a+, Itm2a+); 6) CD4 single positive (SP) cells (Cd4+, Cd8a-, Klf2+, Zbtb7b+); 7) CD8 SP cells (Cd4, Cd8a+, Klf2+, Runx3+).

#### Statistical analysis

Cell-level thymocyte subtype composition was summarized, and donor-level difference among age groups was tested by Kruskal-Wallis rank sum test.

#### Software

All analysis were performed in R v4.2.1 (Vienna, Austria). Seurat V5^72^ was used for Tabula Muris Senis dataset preprocessing, clustering, and annotation.

### Single-cell RNA-seq analysis of the mouse heart from Tabula Muris Senis

Processed single-cell RNA-seq data of the mouse heart were accessed from the Tabula Muris Senis dataset^2^. To determine transcriptomic differences by sex, we subset the data to male and female matched timepoint only (18 months). Differential gene expression analysis was performed using the Seurat v5.0.3 FindMarkers function^72^. Pathway enrichment was determined on significantly differential genes (adjusted *p* value < 0.05 and log_2_(fold change) > 0.75) using g:Profiler^73^ on pathway terms from the KEGG, Reactome, and WikiPathways databases. KEGG metabolic pathway gene lists were used to calculate metabolic pathway score using the Seurat AddModuleScore function. Cell composition was determined based on cell type annotations from Tabula Muris Senis^2^.

### Bulk RNA-seq analysis of mouse muscle from Tabula Muris Senis

Bulk RNA-seq counts data from Tabula Muris Senis^54^ were accessed from the Gene Expression Omnibus database (GEO accession: GSE132040) and processed using DESeq2 v1.40.2^74^. Extracellular matrix related pathway gene lists were obtained from the Broad Molecular Signatures Database to calculate mean pathway scores using log_2_(normalized counts + 1) values. Linear regression was performed on individual genes and pathway scores versus age in R v4.3.2 to determine significantly correlated genes and pathways with age (*p* < 0.05).

### Spatial RNAseq Analysis

Data is taken from spatial transcriptomic data of mouse thymus over time^43^. Plots show gene expression changes across time points. Gene list is found in SI table 1. *p*-values represent significance testing between old vs. young (cutoff is 5 weeks) via two-sided t-test and are corrected for multiple-hypothesis testing using Benjamini-Hochberg correction.

### Metabolic clock construction

The dataset underwent random partitioning into training and test sets, with the test set comprising 20% of the data. Subsequently, the training dataset was utilized for training 18 models employing 2-fold cross-validation to identify the best-performing model. This selected model underwent training on the raw data, followed by training another model of the same type on principal components of the raw data. The coefficients of these models were then normalized and sorted. Next, new instances of the same model type were trained on incrementally increasing subsets of the highest coefficients. This sequential training process began with a model trained solely on the first feature and continued, incorporating additional features until 50% of all features were included. The assumption underlying this approach is that a subset of features can effectively explain the majority of variance. Features that led to an increase in the model’s performance score were retained, resulting in a final selection of features for further analysis.

## Acknowledgments

This work is supported by Impetus Grants 014746-00001. D.A. is supported by the University of Michigan Postdoctoral Pioneer Program and National Institute of Allergy and Infectious Diseases Training Grant T32-AI007413. G.G. is supported by the National Institute on Aging T32AG052374. R.H.S. is supported by the National Institute on Aging R01AG079806 and the Larry L. Hillblom Foundation 2022-A-010-SUP. B.Y. is supported by the Parker Institute for Cancer Immunotherapy, the V Foundation and Donald E. & Delia B Baxter Foundation. P.J.M. is supported by Impetus Grants 014746-00001, the American Lung Foundation COVID-920798 and the Donald E. and Julia B. Baxter Foundation.

## Author contributions

Conceptualization, P.J.M.; Methodology, S.E.P., D.L., L.Z., M.U. and P.J.M.; Formal Analysis, S.E.P., D.L., S.W., S.L. and W.B.T.; Investigation, S.E.P., D.A., E.E., L.Z., X.S., R.M., S.B.K., D.M., P.S., H.W., A.J.C., G.G. and P.J.M.; Resources, B.K.K., C.A.L. and P.J.M.; Writing – Original Draft, S.E.P. and P.J.M.; Writing – Review and Editing, S.E.P. and P.J.M. with input from all authors; Visualization, S.E.P., D.L., S.W., S.L. and W.B.T.; Supervision, R.H.S., B.Y., B.K.K., C.A.L. and P.J.M.; Project Administration, B.K.K., C.A.L. and P.J.M.; Funding Acquisition, P.J.M.

---

**Figure S1:**
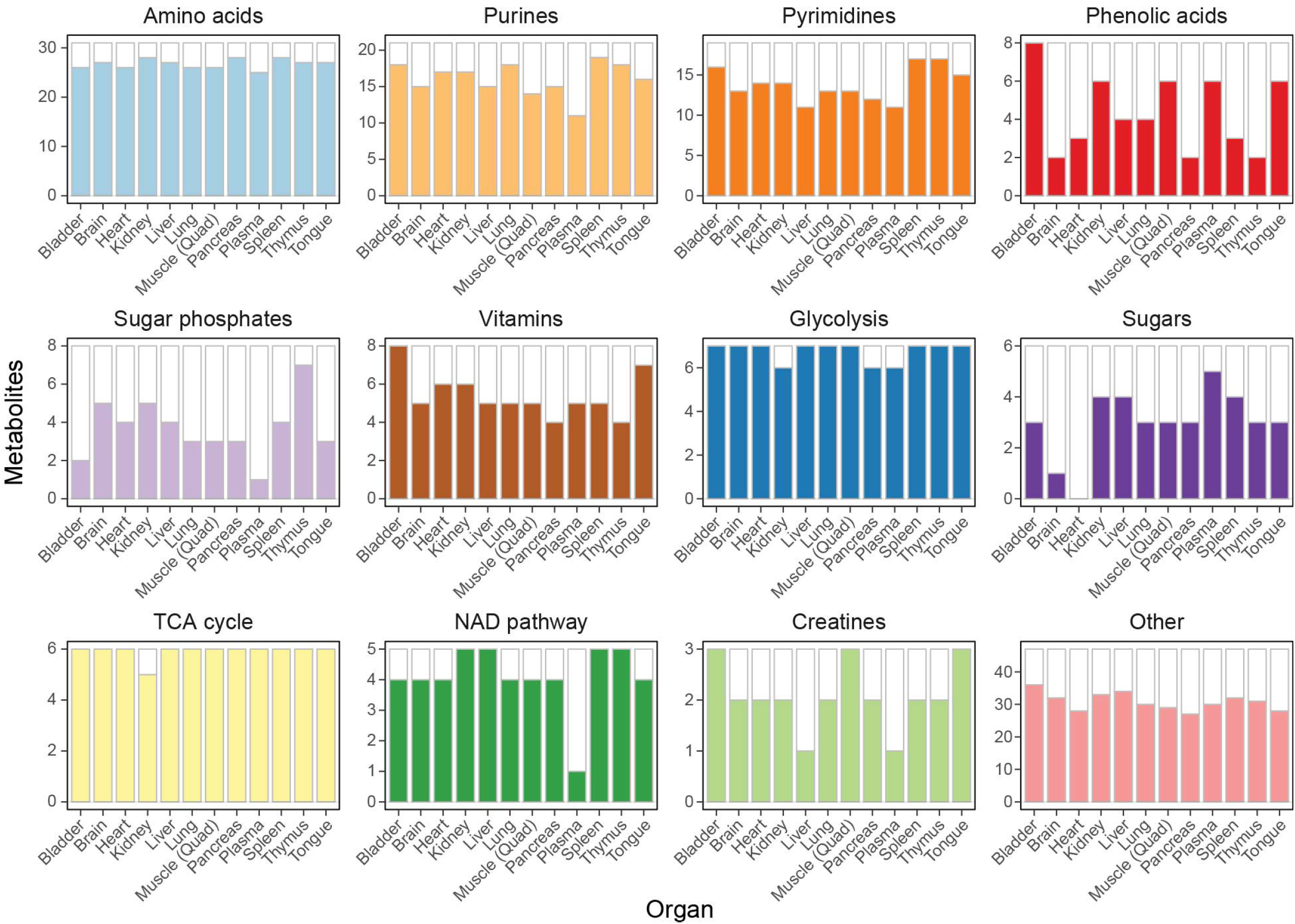
Distribution of metabolite class across organs. Graphs show the number of metabolites detected in each organ for each class of metabolite. Colors indicate the number of metabolites from a group found in that organ.

**Figure S2:**
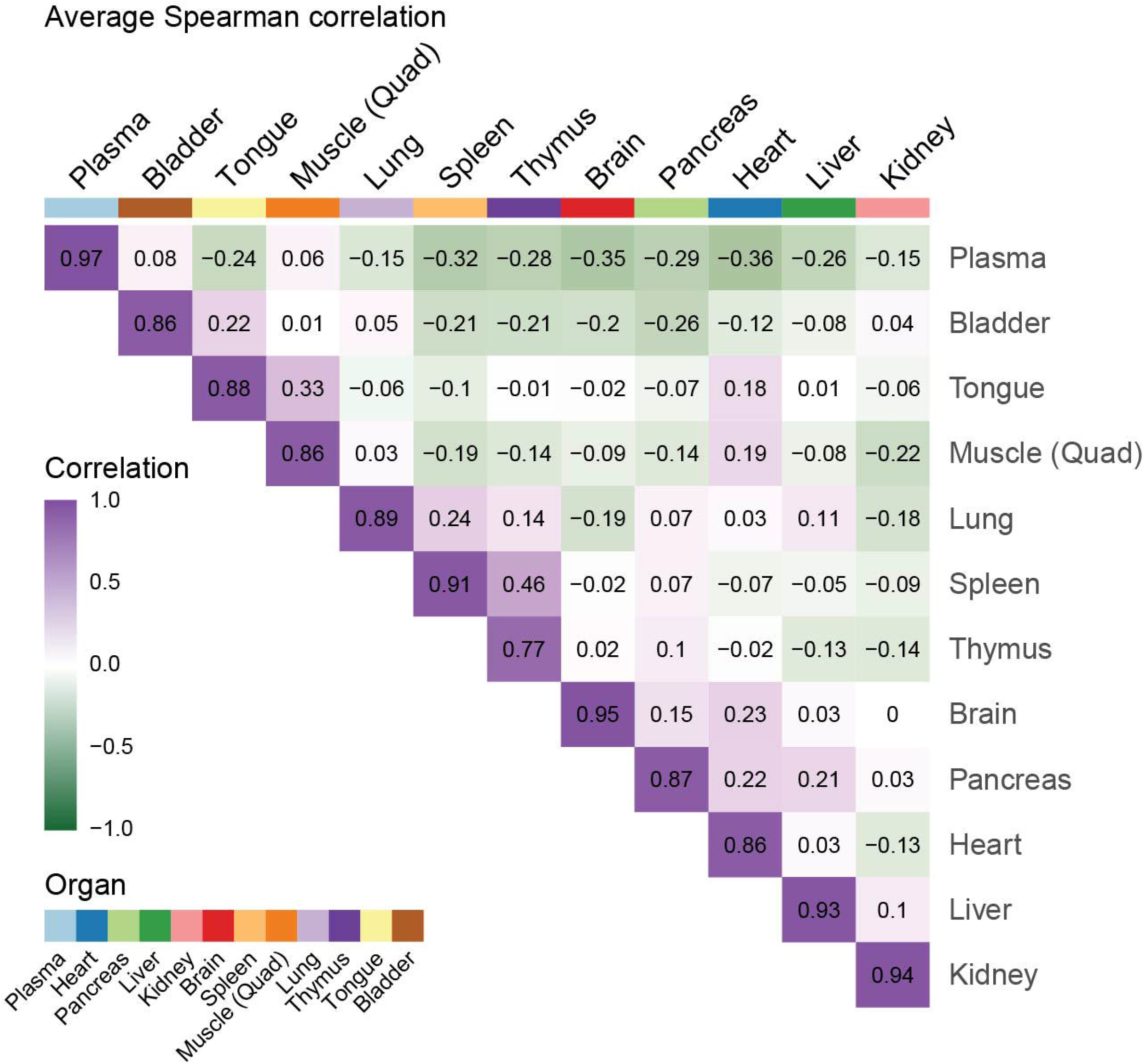
Metabolic correlation varies between organs. Plot showing the mean Spearman correlation of every organ comparison in Figure 2A.

**Figure S3:**
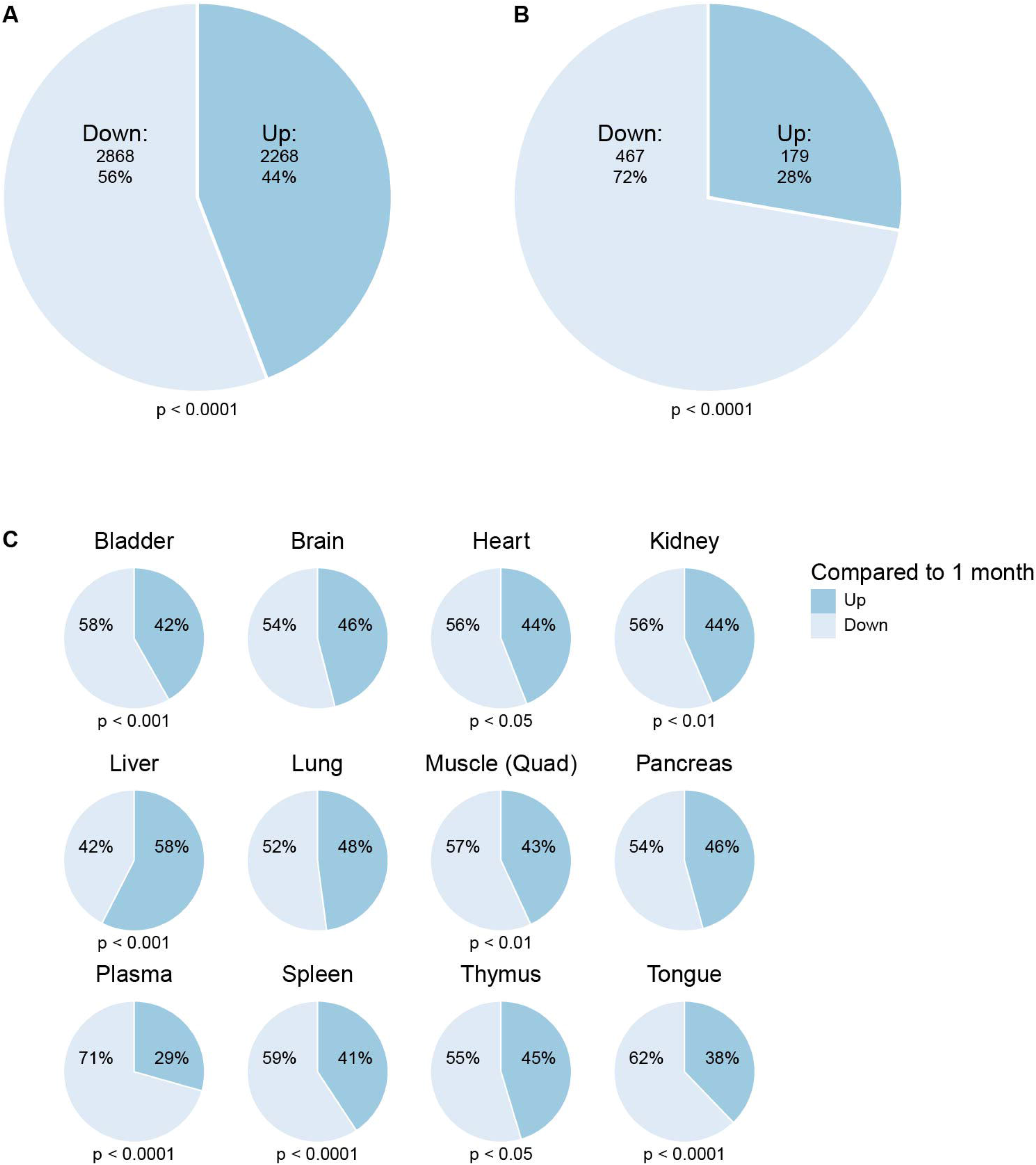
The direction of age-related metabolic changes in individual organs. (A) The majority of comparisons show a decrease with age. Pie chart showing the proportions of total metabolite comparisons that show either an increase (up) or a decrease (down) compared to 1 month. The *p* value was calculated from a binomial test to determine if the proportion of up and down comparisons is significantly different from 50:50. (B) The majority of age-related significant comparisons show a decrease with age. Pie chart showing the proportions of metabolite comparisons which show significant increases (up) or decreases (down) compared to 1 month (FDR < 0.05, log_2_ mean fold change < −1 or > 1). The *p* value was calculated from a binomial test to determine if the proportion of up and down comparisons is significantly different from 50:50. (C) Age-related comparisons show a decrease in 11 out of 12 organs. Pie chart showing the proportions of metabolite comparisons in each organ that show an increase (up) or a decrease (down) compared to 1 month. The *p* values were calculated from a binomial test to determine if the proportion of up and down comparisons is significantly different from 50:50.

**Figure S4:**
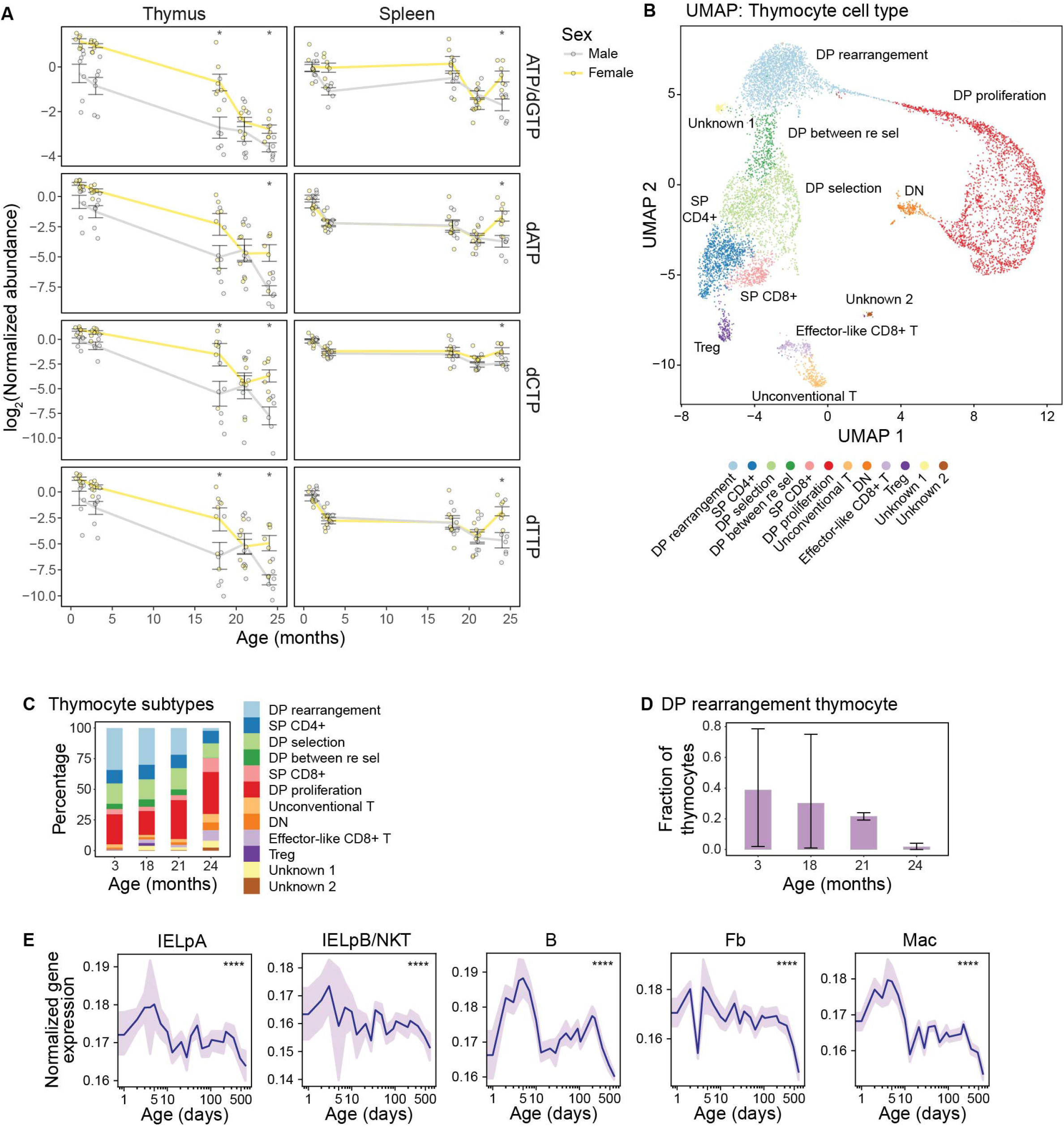
Metabolite and RNA expression data suggest proliferation in the thymus decreases with age. (A) Lines show change in the log_2_ transformed normalized (to 1-month old) abundance of indicated nucleotides in males and females in the thymus and spleen. Error bars show standard error. * FDR < 0.05. (B) UMAP of all thymocytes as found in the Tabula Muris Senis^2^. The analysis used 9338 total thymocytes identified from 9 male mice and 8 female mice. (C) Cellular composition of the thymus at different ages in 9 male mice and 8 female mice. Data analyzed from the Tabula Muris Senis^2^. (D) Fraction of double positive cells undergoing rearrangement from Figure S4C. Error bars show the minimum and maximum values. (E) A gene expression signature of proliferation decreases during aging in multiple cell types. The gene expression signature is shown for the following cells: mouse intraepithelial lymphocyte precursor A (IELpA); mouse intraepithelial lymphocyte precursor B/Natural Killer T (IELpB/NKT); B cell (B); fibroblast (Fb); macrophage (Mac). Significance was calculated between old and young (cutoff at 5 weeks) via two-sided t-test, **** *p* value < 0.0001, corrected for multiple comparisons using Benjamini-Hochberg method.

**Figure S5:**
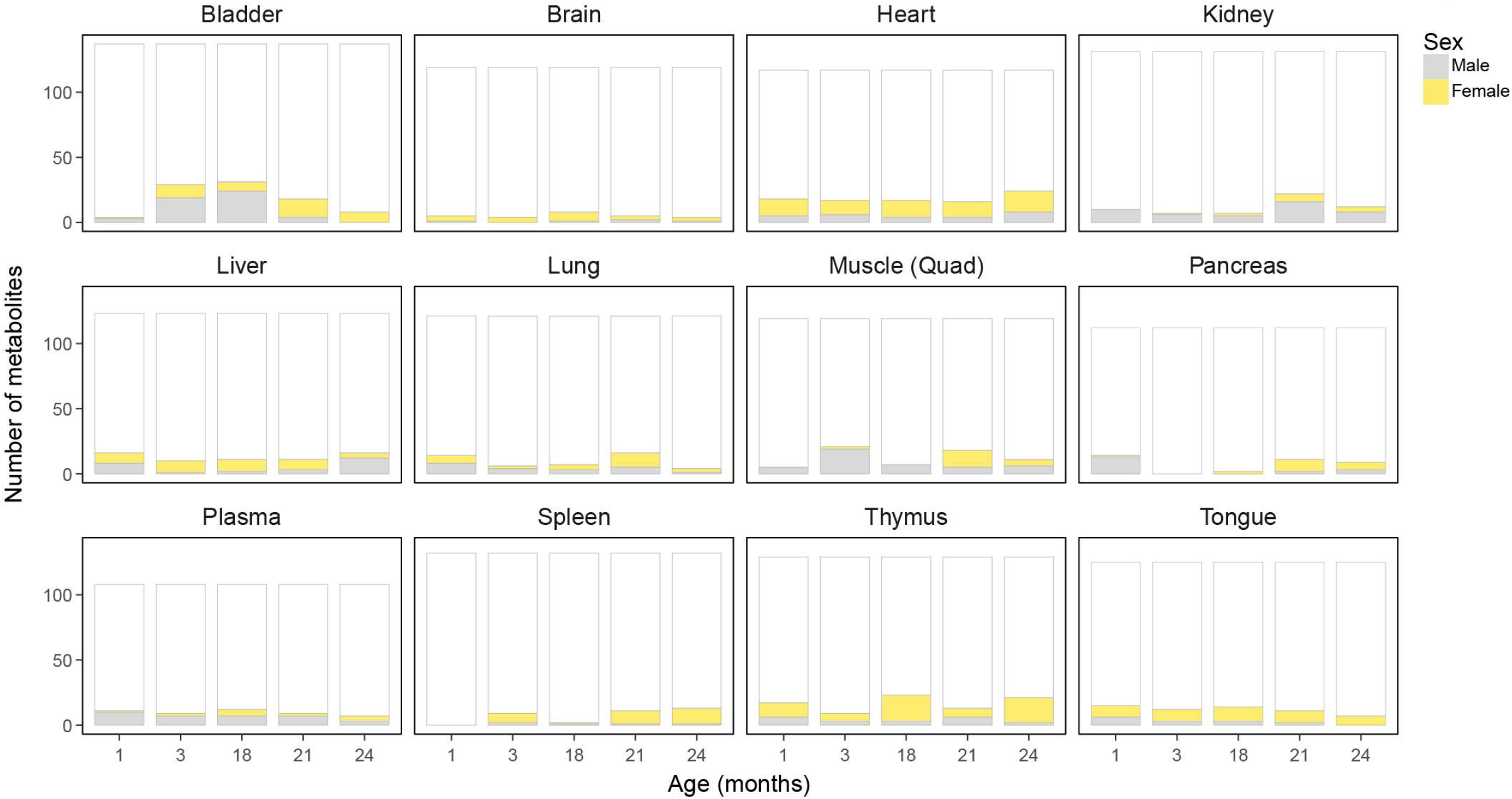
Organs show distinct patterns of metabolic sex differences. **(A)** Bars show the number of metabolites identified in each organ. Shading indicates the number of metabolites that are significantly higher in males compared to females (gray), or higher in females compared to males (yellow), in each age group. White indicates metabolites that show no significant sex difference.

**Figure S6:**
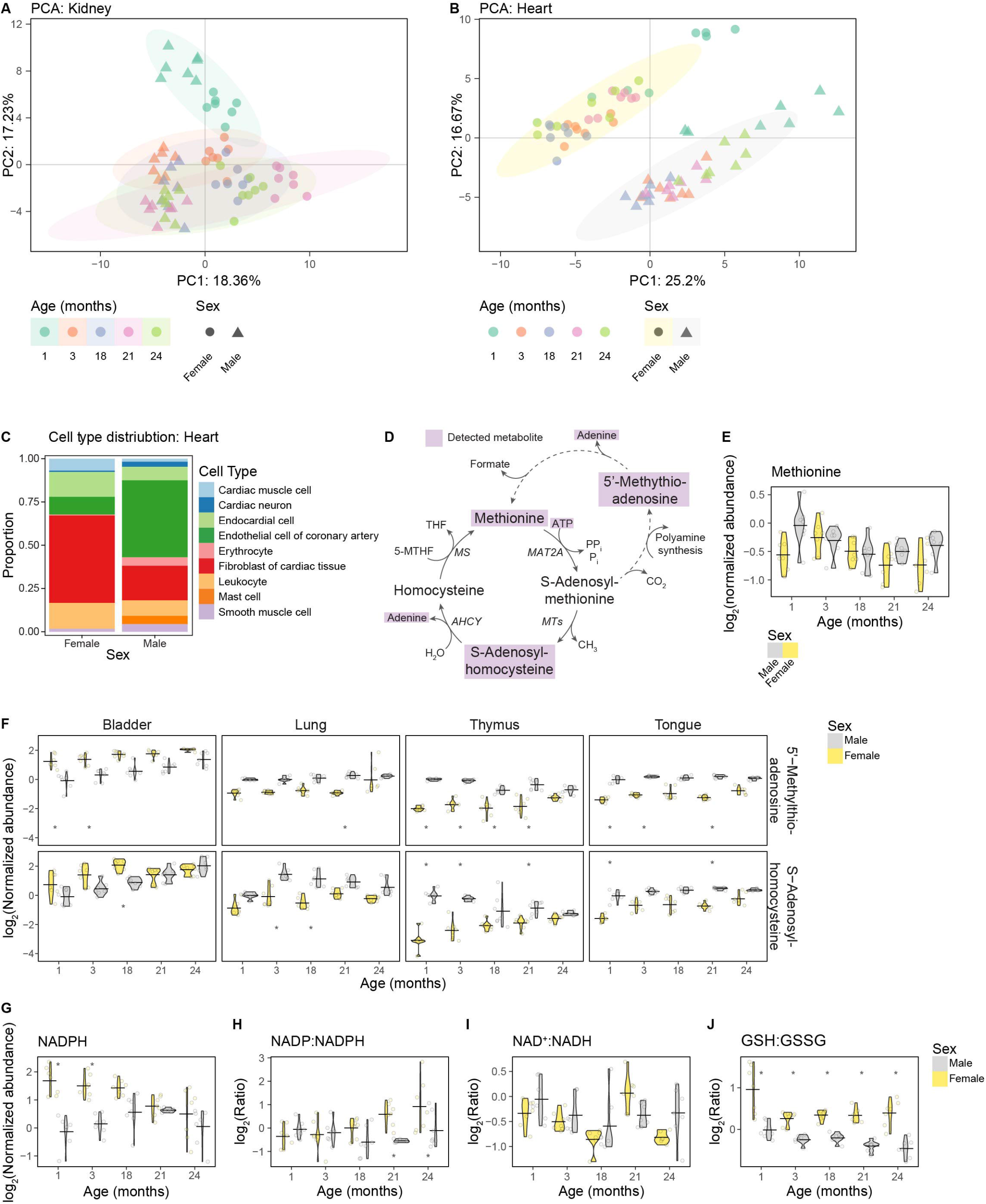
Male and female hearts are metabolically different. (A) Principal component analysis of kidney samples. The shape distinguishes sex and color distinguishes age. (B) Principal component analysis of heart samples. The shape distinguishes sex and color distinguishes age. (C) Cell type distribution in 18-month-old male and female hearts analyzed from single-cell RNA sequencing from the Tabula Muris Senis^2^. (D) Schematic of methionine metabolism. The purple boxes represent metabolites that we identified in our targeted metabolomics analysis. MAT2A = methionine adenosyltransferase 2A, MTs = methyltransferases, AHCY = adenosylhomocysteinase, MS = methionine synthase. (E) Violin plots showing the log_2_ transformed normalized (to 1-month old) abundance of methionine in male and female hearts. Horizontal lines indicate the mean log_2_ transformed normalized abundances at each age. * indicates FDR < 0.05 for male to female comparison. (F) Violin plots showing the log_2_ transformed normalized (to 1-month old) abundances of 5’-methylthioadenosine and S-adenosylhomocysteine in the indicated organs from male and female mice. Horizontal lines indicate the mean log_2_ transformed normalized abundances at each age. * indicates FDR < 0.05 for male to female comparisons. (G) Violin plots showing the log_2_ transformed normalized (to 1-month old) abundance of NADPH in male and female hearts at each age. Horizontal lines indicate the mean log_2_ transformed normalized abundances at each age. * indicates FDR < 0.05 for male to female comparisons. (H) Violin plots showing the log2 transformed NADP:NADPH ratios in male and female hearts at age. * indicates FDR < 0.05 for male to female comparison. (I) Violin plots showing the log2 transformed NAD^+^:NADH ratios in male and female hearts at each age. * indicates FDR < 0.05 for male to female comparison. (J) Violin plots showing the log2 transformed GSH:GSSG ratios in the heart at age. * indicates FDR < 0.05 for male to female comparison.

**Figure S7:**
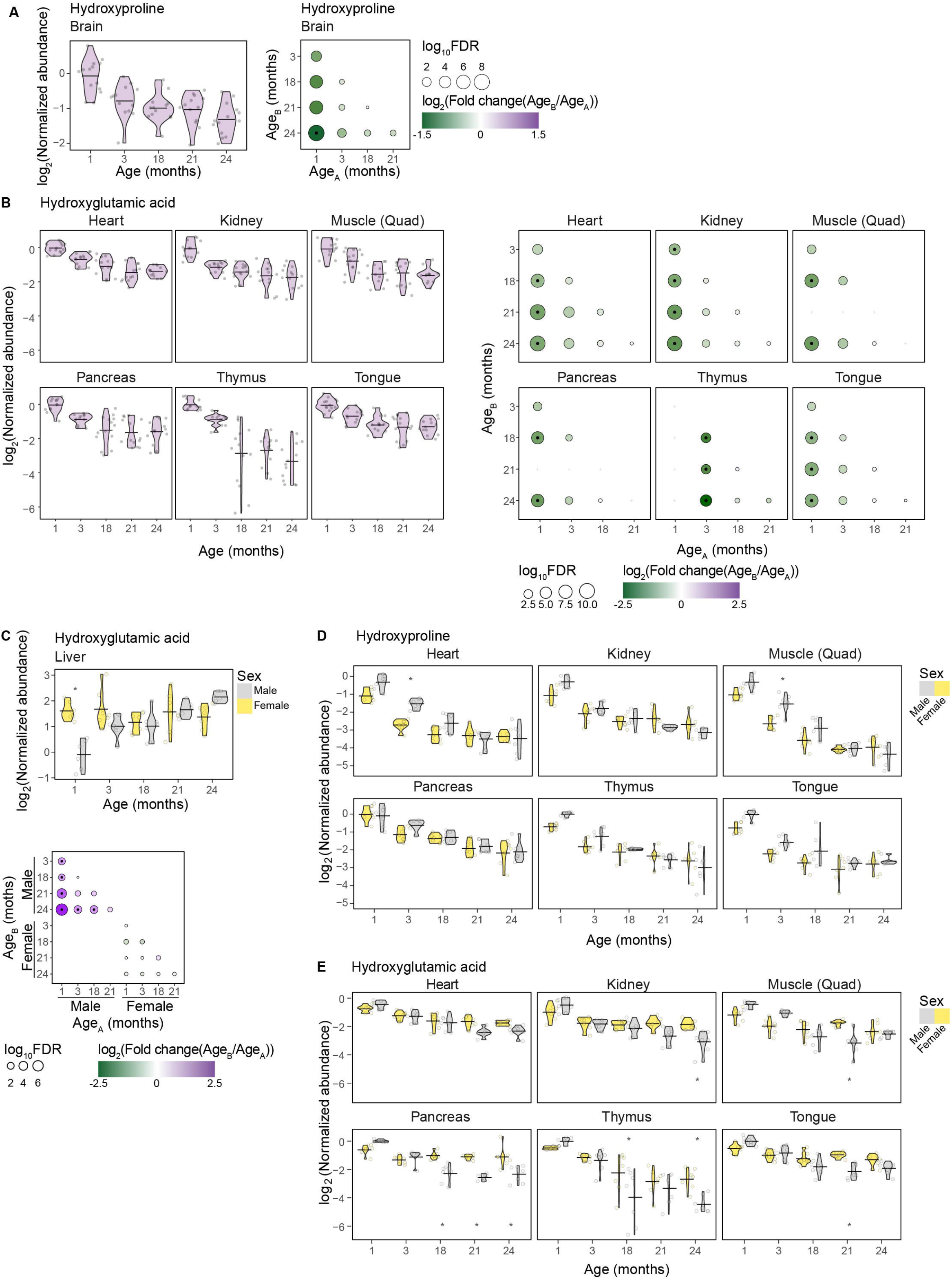
Hydroxyglutamic acid levels decrease with age. (A) Left. Violin plots showing the log_2_ transformed normalized (to 1-month old) abundance of hydroxyproline in the brain. Horizontal lines indicate the mean log_2_ transformed normalized abundances at each age. Right. Significance plots of age-related changes in alpha-ketoglutarate levels. Circle diameter shows the log_10_ FDR for each comparison of age B relative to age A. Circle color indicates the size of the log_2_ transformed fold change. A black dot in the center of the circle indicates the comparison has an FDR < 0.05 and a log_2_ mean fold change of < −1 or > 1. A gray cross indicates the comparison was excluded as the data was found to not be normally distributed. (B) Left. Violin plots showing the log_2_ transformed normalized (to 1-month old) abundance of hydroxyglutamic acid in the indicated organs. Horizontal lines indicate the mean log_2_ transformed normalized abundances at each age. Right. Significance plots of age-related changes in hydroxyglutamic acid levels. Circle diameter shows the log_10_ FDR for each comparison of age B relative to age A. Circle color indicates the size of the log_2_ transformed fold change. A black dot in the center of the circle indicates the comparison has an FDR < 0.05 and a log_2_ mean fold change of < −1 or > 1. A gray cross indicates the comparison was excluded as the data was found to not be normally distributed. (C) Hydroxyglutamic acid levels increase in male livers during aging. Top. Violin plots showing the log_2_ transformed normalized (to 1-month old) abundance of hydroxyglutamic acid in the liver separated by sex. Horizontal lines indicate the mean log_2_ transformed normalized abundances at each age for males and females. Bottom. Significance plots of age-related changes in hydroxyglutamic acid levels. Circle diameter shows the log_10_ FDR for each comparison of age B relative to age A. Circle color indicates the size of the log_2_ transformed fold change. A black dot in the center of the circle indicates the comparison has an FDR < 0.05 and a log_2_ mean fold change of < −1 or > 1. A gray cross indicates the comparison was excluded as the data was found to not be normally distributed.4-hydroxy-L-glutamic acid in the liver. (D) Hydroxyproline levels decrease in multiple organs in males and females during aging. Top. Violin plots showing the log_2_ transformed normalized (to 1-month old) abundance of hydroxyproline in the indicated organs separated by sex. Horizontal lines indicate the mean log_2_ transformed normalized abundances at each age for males and females. Bottom. Significance plots of age-related changes in hydroxyproline levels. Circle diameter shows the log_10_ FDR for each comparison of age B relative to age A. Circle color indicates the size of the log_2_ transformed fold change. A black dot in the center of the circle indicates the comparison has an FDR < 0.05 and a log_2_ mean fold change of < −1 or > 1. A gray cross indicates the comparison was excluded as the data was found to not be normally distributed. (E) Hydroxyglutamic acid levels decrease in multiple organs in males and females during aging. Top. Violin plots showing the log_2_ transformed normalized (to 1-month old) abundance of hydroxyglutamic acid in the indicated organs separated by sex. Horizontal lines indicate the mean log_2_ transformed normalized abundances at each age for males and females. Bottom. Significance plots of age-related changes in hydroxyglutamic acid levels. Circle diameter shows the log_10_ FDR for each comparison of age B relative to age A. Circle color indicates the size of the log_2_ transformed fold change. A black dot in the center of the circle indicates the comparison has an FDR < 0.05 and a log_2_ mean fold change of < −1 or > 1. A gray cross indicates the comparison was excluded as the data was found to not be normally distributed.

**Figure S8:**
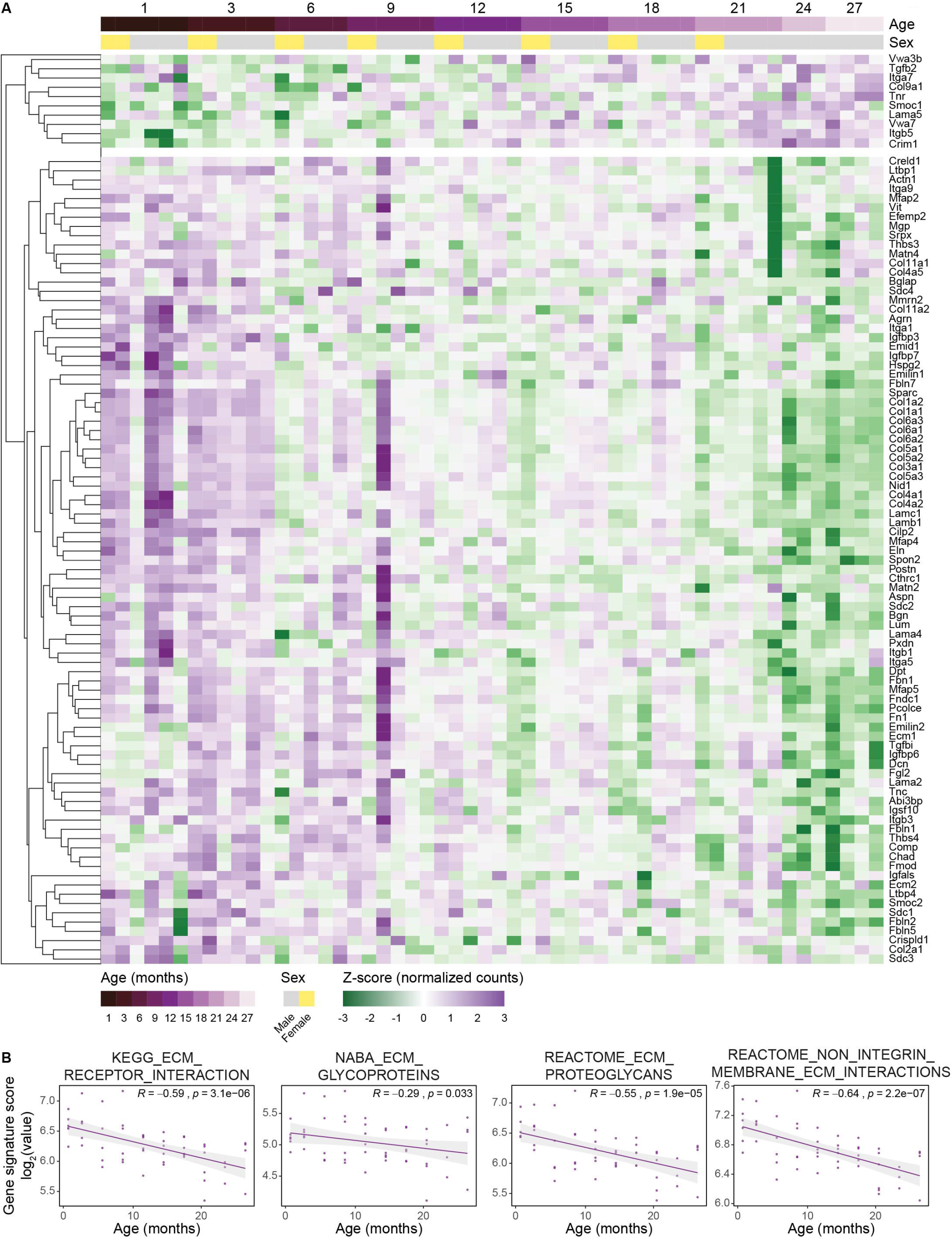
Extracellular matrix gene expression is reduced in quadriceps muscle during aging. (A) Heatmap showing significant changes in the expression ECM-related genes form quadriceps muscle during aging. Data analyzed from the Tabula Muris Senis^2^. (B) Pathway analysis of age-related gene expression changes in the quadriceps muscle from the Tabula Muris Senis^2^. ‘r’ indicates Pearson correlation coefficient.

**Figure S9:**
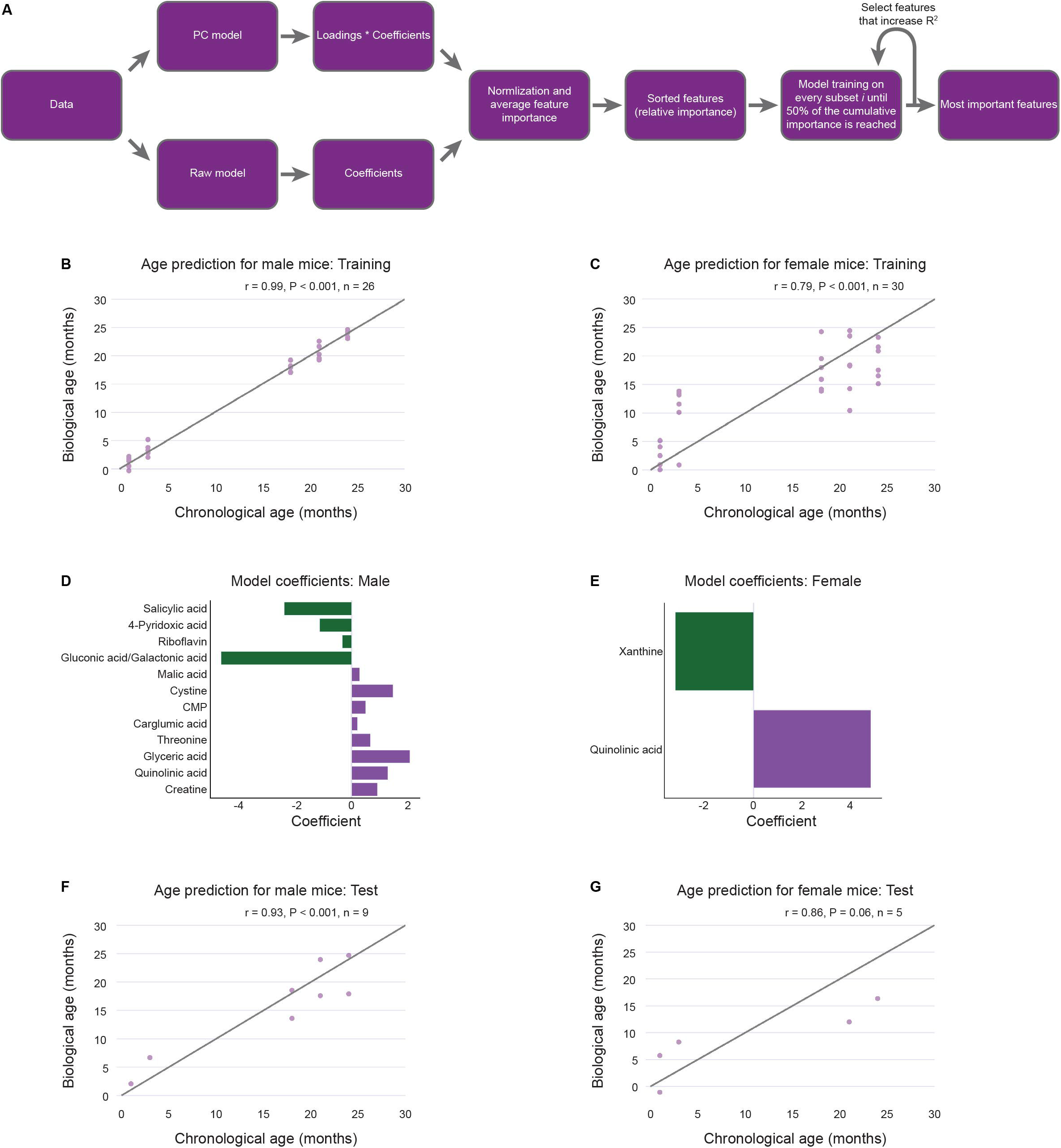
Sex-specific metabolic clocks can also predict age. (A) An overview of how features were selected to build the metabolic aging clocks. (B) Male training set data. The chronological age of male samples plotted against the predicted biological age of male samples in the training set. ‘r’ indicates Pearson correlation coefficient. (C) Female training set data. The chronological age of female samples plotted against the predicted biological age of female samples in the training set. ‘r’ indicates Pearson correlation coefficient. (D) Coefficients of the 12 metabolites used by the model to predict age in the male clock. (E) Coefficients of the 2 metabolites used by the model to predict age in the female clock. (F) Male training set data. The chronological age of male samples plotted against the predicted biological age of male samples in the test set. ‘r’ indicates Pearson correlation coefficient. (G) Female training set data. The chronological age of female samples plotted against the predicted biological age of female samples in the test set. ‘r’ indicates Pearson correlation coefficient.

## References

1. Green, C.L., Lamming, D.W., and Fontana, L. (2022). Molecular mechanisms of dietary restriction promoting health and longevity. Nat Rev Mol Cell Biol 23, 56–73. 10.1038/s41580-021-00411-4.

2. Almanzar, N., Antony, J., Baghel, A.S., Bakerman, I., Bansal, I., Barres, B.A., Beachy, P.A., Berdnik, D., Bilen, B., Brownfield, D., et al. (2020). A single-cell transcriptomic atlas characterizes ageing tissues in the mouse. Nature 583, 590–595. 10.1038/s41586-020-2496-1.

3. Schaum, N., Karkanias, J., Neff, N.F., May, A.P., Quake, S.R., Wyss-Coray, T., Darmanis, S., Batson, J., Botvinnik, O., Chen, M.B., et al. (2018). Single-cell transcriptomics of 20 mouse organs creates a Tabula Muris. Nature 562, 367–372. 10.1038/s41586-018-0590-4.

4. López-Otín, C., Blasco, M.A., Partridge, L., Serrano, M., and Kroemer, G. (2023). Hallmarks of aging: An expanding universe. Cell 186, 243–278. 10.1016/j.cell.2022.11.001.

5. Ikeno, Y., Hubbard, G.B., Lee, S., Cortez, L.A., Lew, C.M., Webb, C.R., Berryman, D.E., List, E.O., Kopchick, J.J., and Bartke, A. (2009). Reduced incidence and delayed occurrence of fatal neoplastic diseases in growth hormone receptor/binding protein knockout mice. Journals of Gerontology - Series A Biological Sciences and Medical Sciences 64, 522–529. 10.1093/gerona/glp017.

6. Miller, R.A., Buehner, G., Chang, Y., Harper, J.M., Sigler, R., and Smith-Wheelock, M. (2005). Methionine-deficient diet extends mouse lifespan, slows immune and lens aging, alters glucose, T4, IGF-I and insulin levels, and increases hepatocyte MIF levels and stress resistance. Aging Cell 4, 119–125. 10.1111/j.1474-9726.2005.00152.x.

7. Gyanwali, B., Lim, Z.X., Soh, J., Lim, C., Guan, S.P., Goh, J., Maier, A.B., and Kennedy, B.K. (2022). Alpha-Ketoglutarate dietary supplementation to improve health in humans. Trends in Endocrinology & Metabolism 33, 136–146. 10.1016/j.tem.2021.11.003.

8. Arriola Apelo, S.I., and Lamming, D.W. (2016). Rapamycin: An InhibiTOR of Aging Emerges From the Soil of Easter Island. J Gerontol A Biol Sci Med Sci 71, 841–849. 10.1093/gerona/glw090.

9. Vermeij, W.P., Hoeijmakers, J.H.J., and Pothof, J. (2016). Genome Integrity in Aging: Human Syndromes, Mouse Models, and Therapeutic Options. Annu Rev Pharmacol Toxicol 56, 427–445. 10.1146/annurev-pharmtox-010814-124316.

10. Hanahan, D., and Weinberg, R.A. (2011). Hallmarks of cancer: the next generation. Cell 144, 646–674. 10.1016/j.cell.2011.02.013.

11. Pilley, S.E., Esparza, E., and Mullen, P.J. (2023). The aging tumor metabolic microenvironment. Curr Opin Biotechnol 84, 102995. 10.1016/j.copbio.2023.102995.

12. Krall, A.S., Mullen, P.J., Surjono, F., Momcilovic, M., Schmid, E.W., Halbrook, C.J., Thambundit, A., Mittelman, S.D., Lyssiotis, C.A., Shackelford, D.B., et al. (2021). Asparagine couples mitochondrial respiration to ATF4 activity and tumor growth. Cell Metab 33, 1013–1026.e6. 10.1016/j.cmet.2021.02.001.

13. Christoffersen, B.Ø., Sanchez-Delgado, G., John, L.M., Ryan, D.H., Raun, K., and Ravussin, E. (2022). Beyond appetite regulation: Targeting energy expenditure, fat oxidation, and lean mass preservation for sustainable weight loss. Obesity 30, 841–857. 10.1002/oby.23374.

14. Lopaschuk, G.D., Karwi, Q.G., Tian, R., Wende, A.R., and Abel, E.D. (2021). Cardiac Energy Metabolism in Heart Failure. Circ Res 128, 1487–1513. 10.1161/CIRCRESAHA.121.318241.

15. Gomes, A.P., Ilter, D., Low, V., Endress, J.E., Fernández-García, J., Rosenzweig, A., Schild, T., Broekaert, D., Ahmed, A., Planque, M., et al. (2020). Age-induced accumulation of methylmalonic acid promotes tumour progression. Nature 585, 283–287. 10.1038/s41586-020-2630-0.

16. Traxler, L., Herdy, J.R., Stefanoni, D., Eichhorner, S., Pelucchi, S., Szücs, A., Santagostino, A., Kim, Y., Agarwal, R.K., Schlachetzki, J.C.M., et al. (2022). Warburg-like metabolic transformation underlies neuronal degeneration in sporadic Alzheimer’s disease. Cell Metab 34, 1248–1263.e6. 10.1016/j.cmet.2022.07.014.

17. Ding, J., Ji, J., Rabow, Z., Shen, T., Folz, J., Brydges, C.R., Fan, S., Lu, X., Mehta, S., Showalter, M.R., et al. (2021). A metabolome atlas of the aging mouse brain. Nat Commun 12. 10.1038/s41467-021-26310-y.

18. Lien, E.C., Vu, N., Westermark, A.M., Danai, L. V, Lau, A.N., Gültekin, Y., Kukurugya, M.A., Bennett, B.D., and Heiden, M.G. Vander (2023). Effects of aging on glucose and lipid metabolism in mice. bioRxiv, 2023.12.17.572088. 10.1101/2023.12.17.572088.

19. Petr, M.A., Alfaras, I., Krawcyzk, M., Bair, W.N., Mitchell, S.J., Morrell, C.H., Studenski, S.A., Price, N.L., Fishbein, K.W., Spencer, R.G., et al. (2021). A cross-sectional study of functional and metabolic changes during aging through the lifespan in male mice. Elife 10. 10.7554/ELIFE.62952.

20. Perez-Ramirez, C.A., Nakano, H., Law, R.C., Matulionis, N., Thompson, J., Pfeiffer, A., Park, J.O., Nakano, A., and Christofk, H.R. (2024). Atlas of fetal metabolism during mid-to-late gestation and diabetic pregnancy. Cell 187, 204–215.e14. 10.1016/j.cell.2023.11.011.

21. Austad, S.N., and Fischer, K.E. (2016). Sex Differences in Lifespan. Cell Metab 23, 1022–1033. 10.1016/j.cmet.2016.05.019.

22. Mauvais-Jarvis, F. (2024). Sex differences in energy metabolism: natural selection, mechanisms and consequences. Nat Rev Nephrol 20, 56–69. 10.1038/s41581-023-00781-2.

23. Clotet-Freixas, S., Zaslaver, O., Kotlyar, M., Pastrello, C., Quaile, A.T., McEvoy, C.M., Saha, A.D., Farkona, S., Boshart, A., Zorcic, K., et al. (2024). Sex differences in kidney metabolism may reflect sex-dependent outcomes in human diabetic kidney disease. Sci Transl Med 16, 13. 10.1126/scitranslmed.abm2090.

24. Horvath, S. (2013). DNA methylation age of human tissues and cell types. Genome Biol 14, R115. 10.1186/gb-2013-14-10-r115.

25. Horvath, S., and Raj, K. (2018). DNA methylation-based biomarkers and the epigenetic clock theory of ageing. Nat Rev Genet 19, 371–384. 10.1038/s41576-018-0004-3.

26. Fitzgerald, K.N., Hodges, R., Hanes, D., Stack, E., Cheishvili, D., Szyf, M., Henkel, J., Twedt, M.W., Giannopoulou, D., Herdell, J., et al. (2021). Potential reversal of epigenetic age using a diet and lifestyle intervention: a pilot randomized clinical trial. Aging 13, 9419–9432. 10.18632/aging.202913.

27. Unfried, M., Ng, L.F., Cazenave-Gassiot, A., Batchu, K.C., Kennedy, B.K., Wenk, M.R., Tolwinski, N., and Gruber, J. (2022). LipidClock: A Lipid-Based Predictor of Biological Age. Frontiers in Aging 3. 10.3389/fragi.2022.828239.

28. Tanaka, T., Biancotto, A., Moaddel, R., Moore, A.Z., Gonzalez-Freire, M., Aon, M.A., Candia, J., Zhang, P., Cheung, F., Fantoni, G., et al. (2018). Plasma proteomic signature of age in healthy humans. Aging Cell 17. 10.1111/acel.12799.

29. Buckley, M.T., Sun, E.D., George, B.M., Liu, L., Schaum, N., Xu, L., Reyes, J.M., Goodell, M.A., Weissman, I.L., Wyss-Coray, T., et al. (2023). Cell-type-specific aging clocks to quantify aging and rejuvenation in neurogenic regions of the brain. Nat Aging 3, 121–137. 10.1038/s43587-022-00335-4.

30. Zhu, H., Chen, J., Liu, K., Gao, L., Wu, H., Ma, L., Zhou, J., Liu, Z., and Han, J.-D.J. (2023). Human PBMC scRNA-seq–based aging clocks reveal ribosome to inflammation balance as a single-cell aging hallmark and super longevity. Sci Adv 9. 10.1126/sciadv.abq7599.

31. Adams, K.J., Pratt, B., Bose, N., Dubois, L.G., St. John-Williams, L., Perrott, K.M., Ky, K., Kapahi, P., Sharma, V., Maccoss, M.J., et al. (2020). Skyline for Small Molecules: A Unifying Software Package for Quantitative Metabolomics. J Proteome Res 19, 1447–1458. 10.1021/acs.jproteome.9b00640.

32. Rother, P., Wohlgemuth, B., Wolff, W., and Rebentrost, I. (2002). Morphometrically observable aging changes in the human tongue. Annals of Anatomy - Anatomischer Anzeiger 184, 159–164. 10.1016/S0940-9602(02)80011-5.

33. Palmer, A.K., and Jensen, M.D. (2022). Metabolic changes in aging humans: current evidence and therapeutic strategies. Journal of Clinical Investigation 132. 10.1172/JCI158451.

34. Menees, K.B., Earls, R.H., Chung, J., Jernigan, J., Filipov, N.M., Carpenter, J.M., and Lee, J.K. (2021). Sex- and age-dependent alterations of splenic immune cell profile and NK cell phenotypes and function in C57BL/6J mice. Immunity and Ageing 18. 10.1186/s12979-021-00214-3.

35. Kaneko, J., Sugawara, Y., Matsui, Y., and Makuuchi, M. (2008). Spleen size of live donors for liver transplantation. Surgical and Radiologic Anatomy 30, 515–518. 10.1007/s00276-008-0364-z.

36. Aspinall, R., Andrew, D., London, R., and 9nh, S. (2001). Gender-Related Differences in the Rates of Age Associated Thymic Atrophy.

37. Gui, J., Mustachio, L.M., Su, D.-M., and Craig, R.W. (2012). Thymus Size and Age-related Thymic Involution: Early Programming, Sexual Dimorphism, Progenitors and Stroma. Aging Dis 3, 280–290.

38. Mittelbrunn, M., and Kroemer, G. (2021). Hallmarks of T cell aging. Nat Immunol 22, 687–698. 10.1038/s41590-021-00927-z.

39. Haynes, B.F., Markert, M.L., Sempowski, G.D., Patel, D.D., and Hale, L.P. (2000). The Role of the Thymus in Immune Reconstitution in Aging, Bone Marrow Transplantation, and HIV-1 Infection. Annu Rev Immunol 18, 529–560. 10.1146/annurev.immunol.18.1.529.

40. Ki, S., Park, D., Selden, H.J., Seita, J., Chung, H., Kim, J., Iyer, V.R., and Ehrlich, L.I.R. (2014). Global transcriptional profiling reveals distinct functions of thymic stromal subsets and age-related changes during thymic involution. Cell Rep 9, 402–415. 10.1016/j.celrep.2014.08.070.

41. Gray, D.H.D., Seach, N., Ueno, T., Milton, M.K., Liston, A., Lew, A.M., Goodnow, C.C., and Boyd, R.L. (2006). Developmental kinetics, turnover, and stimulatory capacity of thymic epithelial cells. Blood 108, 3777–3785. 10.1182/blood-2006-02-004531.

42. Cowan, J.E., Malin, J., Zhao, Y., Seedhom, M.O., Harly, C., Ohigashi, I., Kelly, M., Takahama, Y., Yewdell, J.W., Cam, M., et al. (2019). Myc controls a distinct transcriptional program in fetal thymic epithelial cells that determines thymus growth. Nat Commun 10. 10.1038/s41467-019-13465-y.

43. Liu, S., Friedrich, M.J., Raichur, R., Gong, D., Kehl, N., Gong, Q., Chen, B., Gustafsson, K., Ma, L.L., Scadden, D.T., et al. (2024). Immune decline in aging linked to shifts in thymic architecture. In Review.

44. Sanderson, S.M., Gao, X., Dai, Z., and Locasale, J.W. (2019). Methionine metabolism in health and cancer: a nexus of diet and precision medicine. Nat Rev Cancer 19, 625–637. 10.1038/s41568-019-0187-8.

45. Forney, L.A., Stone, K.P., Gibson, A.N., Vick, A.M., Sims, L.C., Fang, H., and Gettys, T.W. (2020). Sexually Dimorphic Effects of Dietary Methionine Restriction are Dependent on Age when the Diet is Introduced. Obesity 28, 581–589. 10.1002/oby.22721.

46. Casin, K.M., and Kohr, M.J. (2020). An emerging perspective on sex differences: Intersecting S-nitrosothiol and aldehyde signaling in the heart. Redox Biol 31, 101441. 10.1016/j.redox.2020.101441.

47. O’Neill, L.A.J., and Artyomov, M.N. (2019). Itaconate: the poster child of metabolic reprogramming in macrophage function. Nat Rev Immunol 19, 273–281. 10.1038/s41577-019-0128-5.

48. Phang, J.M., Liu, W., and Zabirnyk, O. (2010). Proline metabolism and microenvironmental stress. Annu Rev Nutr 30, 441–463. 10.1146/annurev.nutr.012809.104638.

49. Adams, E., and Frank, L. (1980). Metabolism of Proline and the Hydroxyprolines. Annu Rev Biochem 49, 1005–1061. 10.1146/annurev.bi.49.070180.005041.

50. Phang, J.M. (2023). The regulatory mechanisms of proline and hydroxyproline metabolism: Recent advances in perspective. Front Oncol 12. 10.3389/fonc.2022.1118675.

51. Selman, M., and Pardo, A. (2021). Fibroageing: An ageing pathological feature driven by dysregulated extracellular matrix-cell mechanobiology. Ageing Res Rev 70, 101393. 10.1016/j.arr.2021.101393.

52. Distler, J.H.W., Györfi, A.-H., Ramanujam, M., Whitfield, M.L., Königshoff, M., and Lafyatis, R. (2019). Shared and distinct mechanisms of fibrosis. Nat Rev Rheumatol 15, 705–730. 10.1038/s41584-019-0322-7.

53. Zhao, Y., Zheng, Z., Zhang, Z., Xu, Y., Hillpot, E., Lin, Y.S., Zakusilo, F.T., Lu, J.Y., Ablaeva, J., Biashad, S.A., et al. (2023). Evolution of high-molecular-mass hyaluronic acid is associated with subterranean lifestyle. Nat Commun 14. 10.1038/s41467-023-43623-2.

54. Schaum, N., Lehallier, B., Hahn, O., Pálovics, R., Hosseinzadeh, S., Lee, S.E., Sit, R., Lee, D.P., Losada, P.M., Zardeneta, M.E., et al. (2020). Ageing hallmarks exhibit organ-specific temporal signatures. Nature 583, 596–602. 10.1038/s41586-020-2499-y.

55. Chen, W.J., Lin, I.H., Lee, C.W., and Chen, Y.F. (2021). Aged skeletal muscle retains the ability to remodel extracellular matrix for degradation of collagen deposition after muscle injury. Int J Mol Sci 22, 1–14. 10.3390/ijms22042123.

56. Asadi Shahmirzadi, A., Edgar, D., Liao, C.Y., Hsu, Y.M., Lucanic, M., Asadi Shahmirzadi, A., Wiley, C.D., Gan, G., Kim, D.E., Kasler, H.G., et al. (2020). Alpha-Ketoglutarate, an Endogenous Metabolite, Extends Lifespan and Compresses Morbidity in Aging Mice. Cell Metab 32, 447–456.e6. 10.1016/j.cmet.2020.08.004.

57. Cullins, M.J., and Connor, N.P. (2017). Alterations of intrinsic tongue muscle properties with aging. Muscle Nerve 56, E119–E125. 10.1002/mus.25605.

58. Connor, N.P., Ota, F., Nagai, H., Russell, J.A., and Leverson, G. (2008). Differences in age-related alterations in muscle contraction properties in rat tongue and hindlimb. J Speech Lang Hear Res 51, 818–827. 10.1044/1092-4388(2008/059).

59. Fei, T., Polacco, R.C., Hori, S.E., Molfenter, S.M., Peladeau-Pigeon, M., Tsang, C., and Steele, C.M. (2013). Age-related differences in tongue-palate pressures for strength and swallowing tasks. Dysphagia 28, 575–581. 10.1007/s00455-013-9469-6.

60. Xiong, L., Liu, J., Han, S.Y., Koppitch, K., Guo, J.J., Rommelfanger, M., Miao, Z., Gao, F., Hallgrimsdottir, I.B., Pachter, L., et al. (2023). Direct androgen receptor control of sexually dimorphic gene expression in the mammalian kidney. Dev Cell 58, 2338–2358.e5. 10.1016/j.devcel.2023.08.010.

61. Sadre-Marandi, F., Dahdoul, T., Reed, M.C., and Nijhout, H.F. (2018). Sex differences in hepatic one-carbon metabolism. BMC Syst Biol 12. 10.1186/s12918-018-0621-7.

62. Ignat’eva, N.Y., Danilov, N.A., Averkiev, S. V., Obrezkova, M. V., Lunin, V. V., and Sobol’, E.N. (2007). Determination of hydroxyproline in tissues and the evaluation of the collagen content of the tissues. Journal of Analytical Chemistry 62, 51–57. 10.1134/S106193480701011X.

63. Donald, S.P., Sun, X.Y., Hu, C.A.A., Yu, J., Mei, J.M., Valle, D., and Phang, J.M. (2001). Proline oxidase, encoded by p53-induced gene-6, catalyzes the generation of proline-dependent reactive oxygen species. Cancer Res 61, 1810–1815.

64. Liu, Y., Borchert, G.L., Donald, S.P., Diwan, B.A., Anver, M., and Phang, J.M. (2009). Proline oxidase functions as a mitochondrial tumor suppressor in human cancers. Cancer Res 69, 6414–6422. 10.1158/0008-5472.CAN-09-1223.

65. Pilley, S.E., Hennequart, M., Vandekeere, A., Blagih, J., Legrave, N.M., Fendt, S.-M., Vousden, K.H., and Labuschagne, C.F. (2023). Loss of attachment promotes proline accumulation and excretion in cancer cells. Sci Adv 9. 10.1126/sciadv.adh2023.

66. Olivares, O., Mayers, J.R., Gouirand, V., Torrence, M.E., Gicquel, T., Borge, L., Lac, S., Roques, J., Lavaut, M.N., Berthezène, P., et al. (2017). Collagen-derived proline promotes pancreatic ductal adenocarcinoma cell survival under nutrient limited conditions. Nat Commun 8. 10.1038/ncomms16031.

67. Liu, Y., Mao, C., Wang, M., Liu, N., Ouyang, L., Liu, S., Tang, H., Cao, Y., Liu, S., Wang, X., et al. (2020). Cancer progression is mediated by proline catabolism in non-small cell lung cancer. Oncogene 39, 2358–2376. 10.1038/s41388-019-1151-5.

68. Cooper, S.K., Pandhare, J., Donald, S.P., and Phang, J.M. (2008). A novel function for hydroxyproline oxidase in apoptosis through generation of reactive oxygen species. Journal of Biological Chemistry 283, 10485–10492. 10.1074/jbc.M702181200.

69. Kerk, S.A., Lin, L., Myers, A.L., Sutton, D.J., Andren, A., Sajjakulnukit, P., Zhang, L., Zhang, Y., Jiménez, J.A., Nelson, B.S., et al. (2022). Metabolic Requirement for GOT2 in Pancreatic Cancer Depends on Environmental Context. Elife 11. 10.7554/elife.73245.

70. Butler, A., Hoffman, P., Smibert, P., Papalexi, E., and Satija, R. (2018). Integrating single-cell transcriptomic data across different conditions, technologies, and species. Nat Biotechnol 36, 411–420. 10.1038/nbt.4096.

71. Durinck, S., Spellman, P.T., Birney, E., and Huber, W. (2009). Mapping identifiers for the integration of genomic datasets with the R/ Bioconductor package biomaRt. Nat Protoc 4, 1184–1191. 10.1038/nprot.2009.97.

72. Hao, Y., Stuart, T., Kowalski, M.H., Choudhary, S., Hoffman, P., Hartman, A., Srivastava, A., Molla, G., Madad, S., Fernandez-Granda, C., et al. (2024). Dictionary learning for integrative, multimodal and scalable single-cell analysis. Nat Biotechnol 42, 293–304. 10.1038/s41587-023-01767-y.

73. Kolberg, L., Raudvere, U., Kuzmin, I., Adler, P., Vilo, J., and Peterson, H. (2023). G:Profiler-interoperable web service for functional enrichment analysis and gene identifier mapping (2023 update). Nucleic Acids Res 51, W207–W212. 10.1093/nar/gkad347.

74. Love, M.I., Huber, W., and Anders, S. (2014). Moderated estimation of fold change and dispersion for RNA-seq data with DESeq2. Genome Biol 15. 10.1186/s13059-014-0550-8.

